# Tissue-specific impacts of aging and genetics on gene expression patterns in humans

**DOI:** 10.1101/2021.11.16.468753

**Authors:** Ryo Yamamoto, Ryan Chung, Juan Manuel Vazquez, Huanjie Sheng, Philippa Steinberg, Nilah M Ioannidis, Peter H Sudmant

**Affiliations:** Department of Integrative Biology, University of California, Berkeley; Bioinformatics Interdepartmental Program, University of California, Los Angeles; Center for Computational Biology, University of California, Berkeley; Department of Electrical Engineering and Computer Sciences, University of California, Berkeley

**Author notes:** RY and RC contributed equally to this work.

**Keywords:** aging, genetics, eQTL, Medawar

## Abstract

Age is the primary risk factor for many common human diseases including heart disease, Alzheimer’s dementias, cancers, and diabetes. Determining how and why tissues age differently is key to understanding the onset and progression of such pathologies. Here, we set out to quantify the relative contributions of genetics and aging to gene expression patterns from data collected across 27 tissues from 948 humans. We show that age impacts the predictive power of expression quantitative trait loci across several tissues. Jointly modelling the contributions of age and genetics to transcript level variation we find that the heritability (*h*^2^) of gene expression is largely consistent among tissues. In contrast, the average contribution of aging to gene expression variance varied by more than 20-fold among tissues with 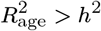 in 5 tissues. We find that the coordinated decline of mitochondrial and translation factors is a widespread signature of aging across tissues. Finally, we show that while in general the force of purifying selection is stronger on genes expressed early in life compared to late in life as predicted by Medawar’s hypothesis, a handful of highly proliferative tissues exhibit the opposite pattern. These *non-Medawarian* tissues exhibit high rates of cancer and age-of-expression associated somatic mutations in cancer. In contrast, gene expression variation that is under genetic control is strongly enriched for genes under relaxed constraint. Together we present a novel framework for predicting gene expression phenotypes from genetics and age and provide insights into the tissue-specific relative contributions of genes and the environment to phenotypes of aging.

## Introduction

Organismal survival requires molecular processes to be carried out with the utmost precision. However, as individuals age many biological processes deteriorate resulting in impaired function and disease. Such increases in the overall variance of molecular processes are predicted by Medawar’s germline mutation accumulation theory (1), which states that because older individuals are less likely to contribute their genetic information to the next generation, there is reduced selection to eliminate deleterious phenotypes that appear late in life (2). This theory also predicts that genes expressed early in life should be under increased selective constraint compared to genes expressed late in life. However, a key challenge remains in both quantifying age-associated changes in biological processes across tissues and identifying how genetic variation influences such changes.

At the organismal level, age-associated changes in the heterogeneity of gene expression between individuals have been observed for a handful of genes in humans (3). In an analysis of gene expression in monozygotic (identical) twins, 42 genes showed age-associated differences in gene expression, suggesting a role for the environment in modulating gene expression with age (2, 3). Similarly, the number of genes with expression quantitative trait loci (eQTLs) detected from blood in 70 year olds declined by 4.7% when they were resampled at 80 years old (4). However, the extent of this phenomenon, both across genes and tissues, remains unclear (5). Age-associated increases in the heterogeneity of gene expression have also been observed at the level of individual cell-to-cell variation; however, only some cell types appear to be impacted (6). In a recent study of immune T-cells from young and aged individuals, no difference in cell-to-cell variability was observed in unstimulated cells, however, upon immune activation the older cells appeared more heterogeneous (7). It is not known why some cell-types and not others may be more likely to exhibit increased cellular variability.

The relationship between the age at which a specific gene is expressed and the force of purifying selection has also recently been explored across a number of species (8, 9). These analyses have broadly confirmed that, on average, genes expressed later in life are under less constraint compared to those expressed early in life. However, how these patterns vary across different tissues and are impacted by genetic variation has not been systematically explored.

Here we set out to understand how aging affects the molecular heterogeneity of gene expression and to model the relative impact of age and genetic variation on this phenotype across tissues. First, using gene expression data from 948 individuals in GTEx V8 (10) we show that age impacts the predictive power of eQTLs, however to varying extents across different tissues and in old and young individuals. Increases in between individual gene expression heterogeneity were associated with these reductions in eQTL power. Using a regularized linear model-based approach to jointly model the impact of both age and genetic variation on gene expression we find that while the average heritability of gene expression is consistent across tissues, the average contribution of age varies substantially. Furthermore, while the genetic regulation of gene expression is similar across tissues, age-associated changes in gene expression are highly tissuespecific in their action. We use this joint model to identify each gene’s age of expression and show that while in most tissues late-expressed genes do tend to be under more relaxed selective constraint, among a handful of highly proliferative tissues the opposite trend holds.

## Results

### Expression quantitative trait loci exhibit varying predictive power in *old* and *young* individuals across several different tissues

To gain insight into how gene regulatory programs might be impacted by aging we analyzed transcriptomic data collected across multiple tissues from 948 humans (GTEx version 8) (10). We hypothesized that aging might dampen the effect of expression quantitative trait loci (eQTLs) due to factors such as increased environmental variance or molecular infidelity (Fig. 1A). To test this hypothesis we first classified individuals into *young* and *old* age groups conservatively grouping individuals above and below the median age (55 years old, Fig. S1), respectively, restricting our analyses to tissues with at least 100 individuals in both groups (27 tissues in total, Fig. S2, Table S1). In each tissue we down-sampled to match the sample size of old and young individuals while additionally controlling for co-factors such as ancestry and technical confounders (methods). Of note, a common approach to controlling for unobserved confounders in large gene expression experiments is to probabilistically infer hidden factors using statistical tools such as PEER (11). We noticed that many of the GTEx PEER factors were significantly correlated with sample age, with the top three correlated PEER factors having a Pearson r of 0.33, -0.21, and -0.15 (Fig. S3). To prevent loss of age related variation, we recalculated a corrected set of PEER factors that were independent of sample age (Methods). We then assessed the significance of GTEx eQTLs in the young and old cohorts respectively, comparing the distribution of P-values over all genes between old and young individuals (Fig. 1A 1B, two-sided Welch’s t-test). In 20 out of 27 (74%) of the assessed tissues, the P-value distribution was significantly different between young and old individuals with genotypes more predictive of expression in younger individuals in 12/20 cases (Fig. 1C). These results were largely identical when the analyses were performed with the original non-corrected PEER factors (18/27 tissues, Fig. S4). While the GTEx dataset is unique in its wide sampling of participant ages and tissues, we validated our observations in the PIVUS cohort which includes blood tissue from individuals re-sampled at ages 70 and 80 (4). This study previously demonstrated a reduction in eQTL heritability with age supporting our results. We confirmed using our approach that eQTLs were less predictive of gene expression in 80, compared to 70 year old’s (Fig.S5, S6). These results suggest that the predictive power of eQTLs is impacted by the sample age across the vast majority of tissues. Furthermore this effect is more pronounced in older samples compared with younger samples.

**Fig. 1.**
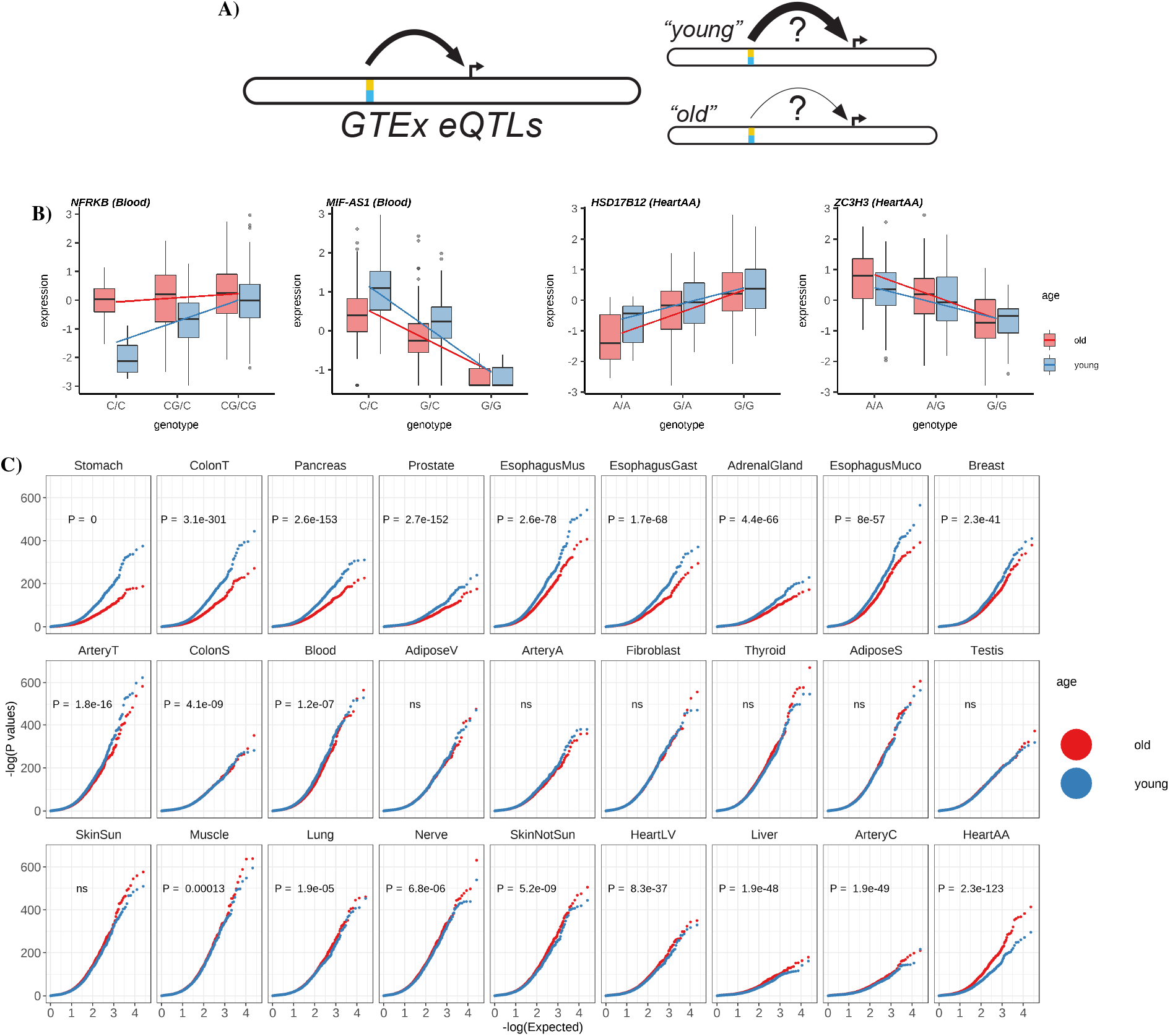
Age impacts the predictive power of eQTLs in many tissues. **A)** A hypothetical model of the differing power to detect eQTLs in old and young cohorts. **B)** Examples of gene expression binned by genotype and age for four genes in which eQTL p-values change in whole blood and heart atrial appendage. Example genes in whole blood show lower p-values within young individuals and lower p-values within old individuals in heart atrial appendage. NFRKB and HSD17B12 show positive effect size while MIF-AS1 and ZC3H3 show negative effect size. Center line of the boxplot indicates median, box limit indicates first and third quartiles and points indicate outliers. **C)** QQ plots of eQTL p-values (plotted as -log(P)) for old (red) and young (blue) individuals from a linear model correlating expression with the lead SNP for each gene in all 27 tissues (Table S2). P-value for significance differences in eQTL p-value distributions are obtained from two-sided Welch’s t-test.

### Age-associated changes in gene expression heterogeneity impact gene expression heritability

We hypothesized that the overall reduced predictive power of eQTLs in some tissues might be in part due to an increase in expression heterogeneity in these tissues, potentially as a result of increased environmental variance. To test if such an effect would broadly affect expression across all genes in a tissue (Fig. 2A) we calculated the distribution of pairwise distances among individual’s tissue-specific gene expression profiles using the Jensen-Shannon Divergence (JSD) (12, 13) as a distance metric. The JSD is a robust distance which is less impacted by outliers compared to other methods (e.g. Euclidean distance) (13). Comparing the distribution of pairwise differences in transcriptional profiles within distinct age groups allows us to determine if gene expression signatures are more similar among *younger* individuals or among *older* individuals.

**Fig. 2.**
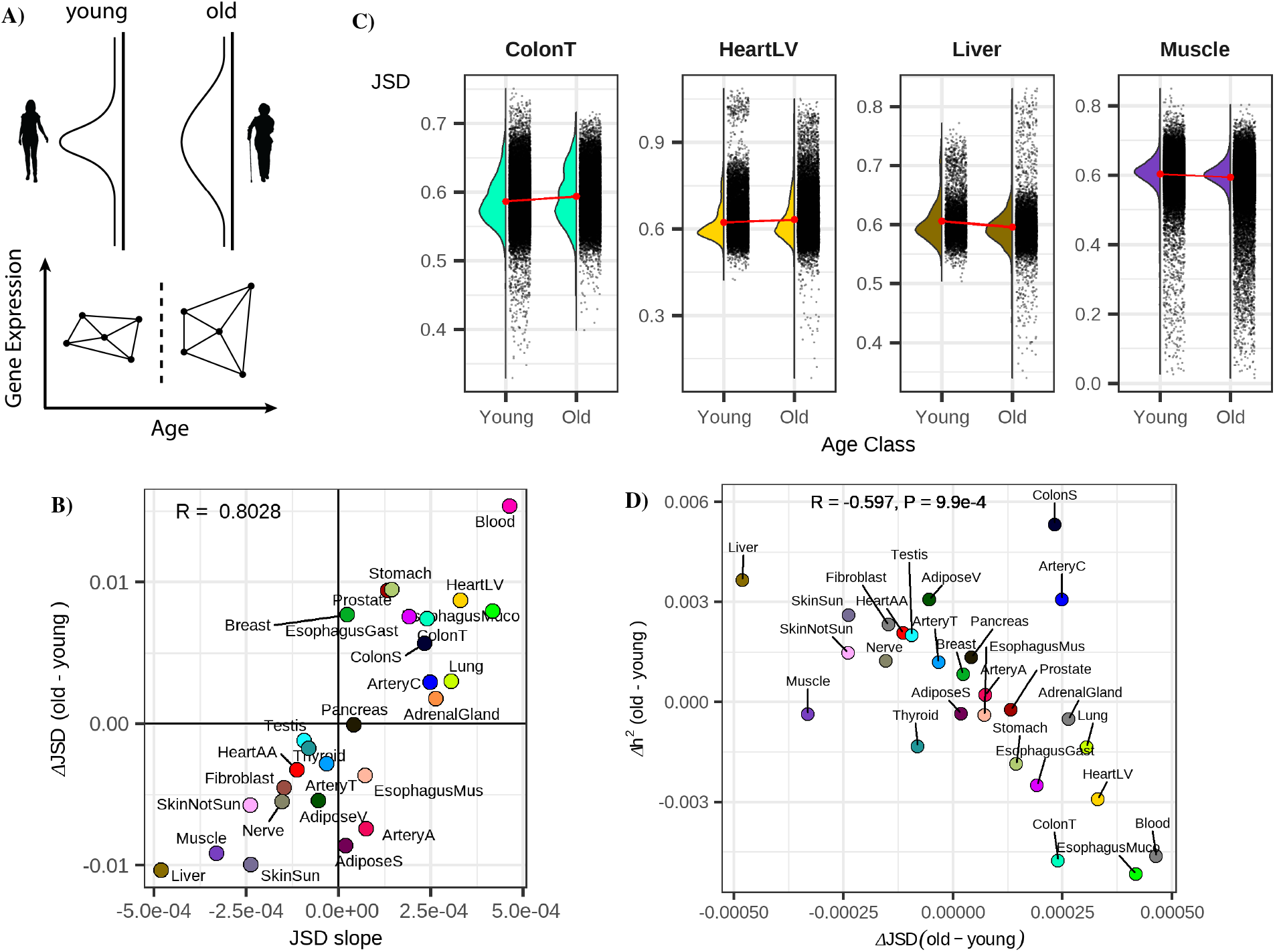
Inter-individual gene expression heterogeneity changes with age for a subset of tissues. **A)** Hypothesized age-associated increase in gene expression heterogeneity (top) and our approach for quantifying the inter-individual expression distance with age using the Jensen Shannon Divergence metric (JSD) for age-binned individuals (bottom). **B)** Consistency of measuring the average age-associated change in gene expression heterogeneity across a tissue using a binary binning strategy (y-axis, *JSD*_*old*_-*JSD*_*young*_) or a 6 bin strategy (x-axis, slope of JSD across 6 bins). R indicates Pearson correlation value **C)** The distributions of JSD distances for four example tissues in old and young bins. **D)** The relationship between gene expression heterogeneity and the difference in expression heritability between old and young individuals. R and p-value are obtained from Pearson correlation test

We compared the mean difference in gene expression distances among old and young individuals as well as the slope of the inter-individual JSD and when grouping individuals into six bins spanning 20-80 years old (see methods, Fig. 2B, 2C). These two strategies yielded highly similar results (Fig. 2C Pearson’s R=0.8) identifying tissues exhibiting increased heterogeneity in both *young* and *old* populations. (Fig. S7) The difference in JSD between old and young individuals was also negatively correlated with the results from our analysis of eQTLs across old and young individuals (Fig. S8, R=-0.48, P=0.01 two-sided Pearson correlation test) highlighting that tissues with increases in inter-individual heterogeneity were likely to also exhibit reductions in the proportion of variance described by eQTLs.

To expand our eQTL analyses to account for the combined impact of nearby SNPs, we utilized the multi-SNP regularized linear model of PrediXcan (14). This model has the benefit of combining genetic effects across many loci, instead of examining just a single eQTL variant. This combined genetic contribution to gene expression variance results in an estimate of the heritability (*h*^2^) for each gene. We applied this model independently in old and young individuals to quantify *h*^2^ and found that the average per-gene difference in *h*^2^ between old and young individuals was strongly negatively correlated with the difference in JSD between samples (Pearson’s R=0.6, P=9.9e-4 Pearson correlation test, Fig. 2D, Fig. S9). To verify these results we again referred to the PIVUS study and obtained cis heritability estimates using the GCTA package (15). As expected, we observed that the heritability of gene expression decreases with age, corresponding with the Predixcan results in GTEx whole blood (Fig. S10). Together these results suggest that across numerous tissues gene expression heterogeneity differs between *young* and *old* individuals. This increased expression variance drives a reduction in the average heritability of gene expression across these tissues.

We additionally sought to identify individual genes exhibiting age-associated expression heterogeneity by testing if, after regressing out age-related changes in gene expression levels, the variance of the residuals correlated with age (*Breusch-Pagan test*). The effect size from this test (*β*_*het*_) describes the strength and direction of age related changes in gene expression variance. Using this approach we identified 279 genes with age-associated variance changes (FDR<0.05) across tissues (Fig S11). The estimated *β*_*het*_ values in these genes were overwhelmingly negative (234/279, 84%, Table S3) indicating that the dominant signature was of reduced gene expression heterogeneity with age. A Gene Set Enrichment Analysis (GSEA) of these genes highlighted pathways involved in metabolism, cell proliferation, cell cycle and cell death (Fig. S12). While the proportion of positively heteroskedastic genes was weakly correlated with the transcriptome-wide JSD (p=1.32e-2 two-sided Pearson correlation test, Fig. S13), the small number of genes implicated suggests that these metrics are capturing different phenomena.

### Cell-type specific age-associated changes in gene expression heterogeneity and the predictive power of eQTLs

While no datasets of the magnitude and scale of GTEx exist for single-cell genomic data, we employed the tool CIBERSORT (16) to deconvolute bulk GTEx blood RNA-seq data into cell-type specific abundances. Assessing the predictive power of eQTLs in *old* and *young* individuals in six immune cell subtypes we again found significantly increased explanatory power of eQTLs in younger individuals compared to older individuals, (Fig. S14). Consistent with these analyses, a comparison of the JSD in *old* and *young* individuals revealed increased expression heterogeneity across these cell types with age (Fig. S15). We also investigated whether the observed differences in eQTL power and expression heterogeneity might be driven by changes in cell-type composition; however, cell-type composition changes were not reflective of gene expression variance (P=0.2 two-sided Pearson correlation test, Fig. S16), suggesting that age associated changes in eQTL power and expression heterogeneity are taking place at the transcript level.

### Jointly modeling the impact of age and genetics on gene expression identifies distinct, tissue-specific patterns of aging

A more powerful approach to understand how both genetics and age impact gene expression variation is to jointly model these factors simultaneously. We set out to extend the regularized linear model employed by PrediX-can (14) to incorporate an age factor (Fig. 3A) allowing us to parse apart the individual contributions of genetics (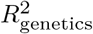 or *h*^2^), age 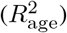, and the environment 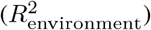, to the expression variance of each gene (e.g. Fig. 3B, Fig. 3C, Fig. S17). We define 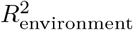 as all sources of variation not captured by *h*^2^ and 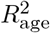. Estimates of *h*^2^ in our extended model were highly consistent with those in the original PrediXcan approach (Fig. S 18).

**Fig. 3.**
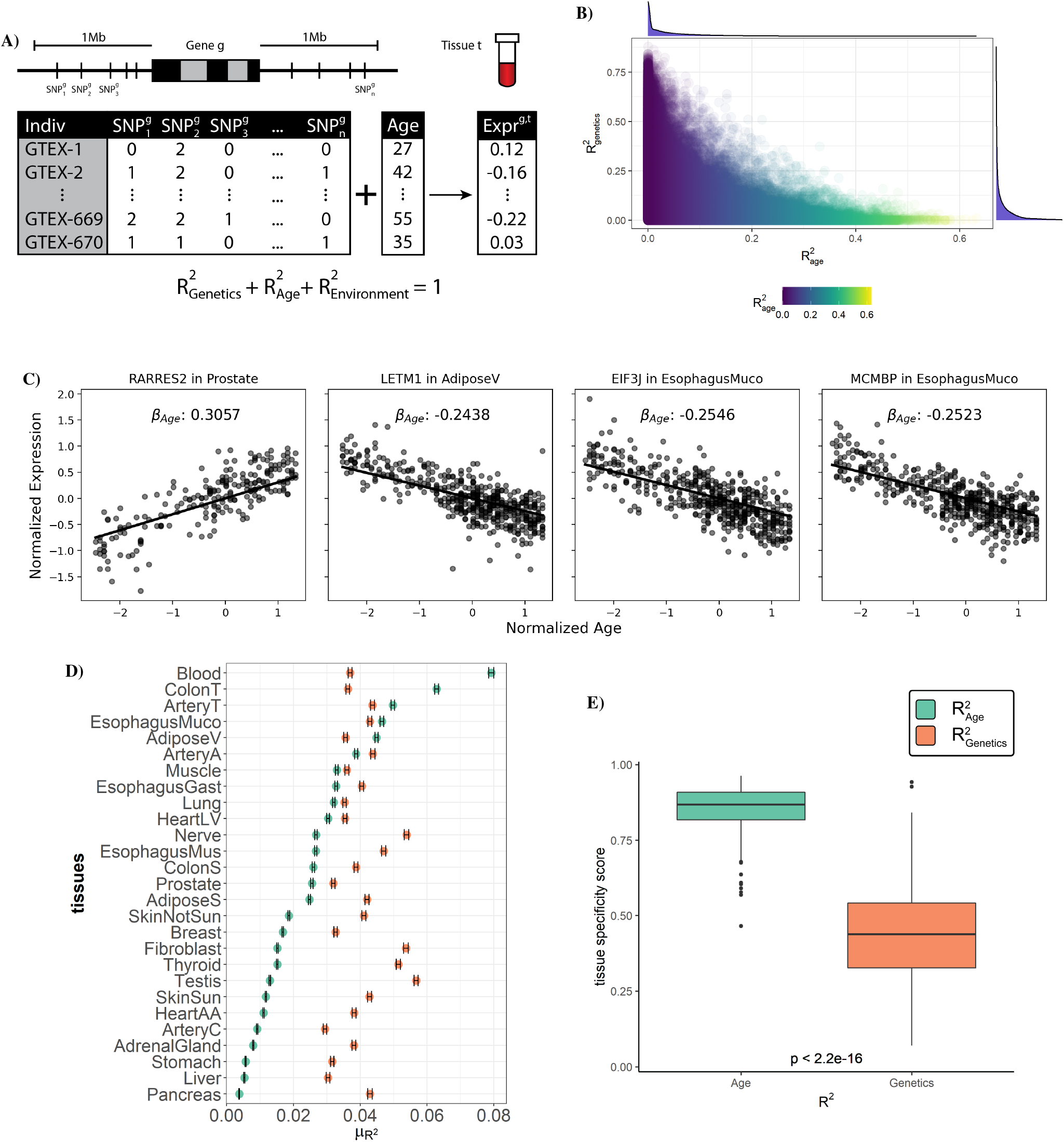
A joint predictive model of gene expression identifies tissue-specific contributions of age and genetics to transcript levels. **A)** A schematic of our multi-SNP gene expression association model incorporating sample age. Common SNPs around each gene *g* are used in combination with an individual’s age to predict expression within tissue *t*. Using this trained model, variation in gene expression can be separated into three parts: the components explained by genetics (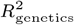 or *h*^2^), by age 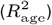 and by all other factors 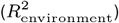. **B)** Proportion of each gene’s expression variance explained by age and genetics. **C)** Plot of normalized expression vs age for four genes with age-correlated expression. Line shows fitted *β*_age_ from regularized linear model. **D)** Point estimates of the mean 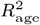 and *h*^2^ for each tissue, error bar indicating confidence interval for the estimate. **E)** The tissue specificity score of *R*^2^ across 27 tissues for each gene from either age genetics. Center line of the boxplot indicates median, box limit indicates first and third quartiles, points indicate outliers. P-value is obtained from two-sided paired samples t-test.

Employing our model across each tissue independently we find that average heritability of gene expression was largely consistent among tissues ranging from 2.9%-5.7% with 40% of genes having an *h*^2^>10% in at least one tissue (Fig. 3D, S19). Thus, while the variation in expression of many individual genes is strongly influenced by genetics, on average, genetics explains a small proportion of overall gene expression variation. In contrast, the average contribution of aging to gene expression varied more than 20-fold among tissues from 0.4%-7.9% with the average 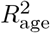 greater than the average *h*^2^ in 5 tissues. Among these 5 tissues the expression of 39-54% of genes was more influenced by age than by genetics (i.e. 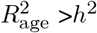, Fig. S20) and across all tissues 45% of genes had an 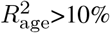 in at least one tissue. Assessing the tissue-specificity of these trends on a per-gene basis we found while the estimated heritability of gene expression tended to be similar among different tissues, the age-associated component exhibited significantly more tissue specificity (P<2.2e-16 two-sided paired t-test, Fig. 3E). We note that the widespread signatures of age-associated gene expression variance that we identify are virtually undetectable when using the GTEx-provided PEER factors. Just 1.84% of the age-associated genes we identify have nonzero age coefficient when using these GTEx PEER factors (Fig. S21). We tested if sex-specific age effects were contributing to the observed age associations as might be expected if changes related to menopause were playing a role (Fig. S22). Including an interaction term between age and sex in our joint model we found that while the age term continued to describe a large proportion of the variance (on average 2.6%), the contribution of the age-sex interaction term was severalfold lower (average of 0.035%, Fig. S23, Table S4). The model incorporating age-sex interactions also showed consistent estimates of variance explained as compared to the baseline joint model (R=0.99 two-sided Pearson correlation test, Fig. S24). Our model thus widely expands the utility of the GTEx dataset and exploration of critical biological signatures of aging. Together these results imply that age-associated patterns of gene expression exhibit substantially more tissue specificity than those that are influenced by genetics and among several tissues age plays a much stronger role in driving gene expression patterns than genetics.

### Coordinated decline of mitochondrial and translation factors is a widespread signature of aging across tissues

To understand the underlying biological implications of age-associated gene expression changes we applied gene set enrichment analysis (GSEA)(17) to each tissue independently, ranking genes either by the relative contribution of genetics (*h*^2^) or aging 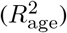. Comparing the distribution of P-values from enriched GO-annotations we found that pathways enriched for age-associated variance were substantially more enriched for significance than pathways associated with genetic-associated variance (e.g. Fig. 4A). We found more age-associated pathway enrichment even in tissues for which the average age-associated contribution to gene expression was low (e.g. Pancreas, Fig. S25). This implies that while age-associated changes in gene expression vary widely in their magnitude among tissues, these changes consistently impact critical biological processes. A GSEA enrichment analysis of genes ranked by the tissue-averaged slope of the age-associated trend (*β*_age_) highlighted several key aging-associated pathways. Pathways associated with various mitochondrial and metabolic processes and translation were enriched for having *−β*_age_ values, implying age-associated decreases (Fig. 4B). A single immune pathway, the interferon-gamma response was enriched having +*β*_age_ values (Fig. 4B). An additional 18 immune pathways were identified as having age-associated increases in gene expression using a more lenient significance threshold (FDR<0.05) (Fig. S26, Table S5). In contrast, no pathways were significantly enriched when genes were ranked by average *h*^2^.

**Fig. 4.**
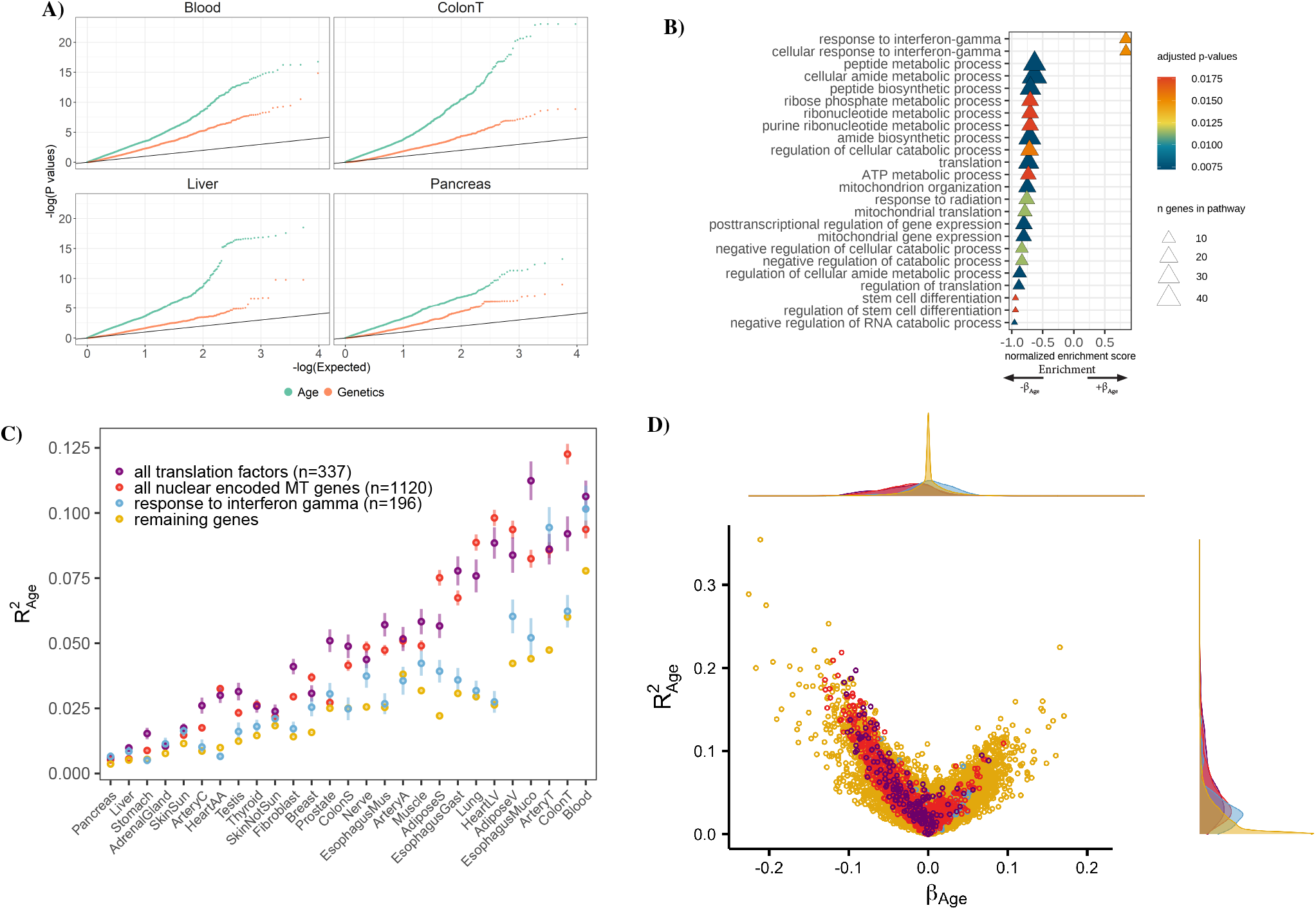
Functional analysis of age-related genes reveals enriched biological processes. **A)** A QQ plot of p-values for pathways tested for enrichment using gene set enrichment analysis (GSEA) with genes ranked by either *h*^2^ or 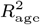 in four example tissues. **B)** GSEA enrichments from genes ranked by the mean *β*_age_ across tissues. Pathways with a correct *P <* 0.02 are shown. **C)** Average gene expression variance explained by age for mitochondrial (MT) genes (red), translation factor genes (purple), interferon gamma genes (blue) and remaining genes (yellow) across all tissues, error bar indicating standard error of point estimates. **D)** Volcano plot of the variance explained by age vs *β*_age_ for mitochondrial, translation factors, interferon gamma factors, and remaining genes. Density plot of each axis show on top and right.

To further explore the functional impact of age-associated gene expression changes we compared the 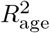 of all nuclear-encoded mitochondrial genes (n=1120, (18)), and translation initiation, elongation, and termination factors, across tissues (Fig. 4C, Fig. S27). Genes in these pathways were exceptionally enriched for age-associated gene expression across several tissues. In some cases >10% of the average expression variation of mitochondrial or translation factor genes could be explained by age. *β*_age_ was consistently negative in these mitochondrial and translation factor genes (Fig. 4D) highlighting that genes in these pathways exhibit a systematic decrease in expression as a function of age. Overall across tissues an average of 36% of all mitochondrial genes (406/1120), and 35% of translation factors (119/337) exhibited age-associated declines, however in some tissues these proportions exceeded 60%. In contrast, the only pathway associated with age-associated increases in expression, interferon-gamma response genes, was largely specific to blood and arterial tissue (Fig. 4C), likely due to the role of this pathway in immune cells. Together these results demonstrate that the coordinated decline of mitochondrial genes and translation factors is a widespread phenomenon of aging across several tissues with potential phenotypic consequences.

### Distinct evolutionary signatures of gene expression patterns influenced by aging and genetics

Evolutionary theory predicts that due to the increased impact of selection in younger individuals, genes that increase as a function of age (*β*_age_ > 0) should be under reduced selective constraint compared to genes that are highly expressed in young individuals (*β*_age_ < 0), a theory of aging known as *Medawar’s hypothesis* (1) (Fig. 5A). Several recent studies have demonstrated the generality of this trend across species (8, 9, 19) however the tissue-specificity of this theoretical prediction has not been explored. We sought to test the generality of this trend across different tissues by comparing *β*_age_ with the level of constraint on genes, quantified as the probability loss of function intolerance (pLI) score from gnomAD (20). As expected, across the vast majority of tissues *β*_age_ was significantly negatively correlated with pLI (Fig. 5B, 5C, S28), in line with Medawar’s hypothesis. However, five tissues exhibited significant signatures in the opposite direction including prostate, transverse colon, breast, whole blood, and lung tissue (P < 10^−3^ linear model two-sided t-test). These five tissues still maintained a significant negative relationship after subsetting to genes that are highly dependent on age (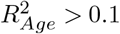, Fig. S29). These tissues with *non-Medawarian* trends are driven by highly constrained, functionally important genes being expressed at a higher rate in older individuals (Fig. S30). Using *dN/dS* (21) as an alternative metric of gene constraint yielded highly correlated results (R=-0.72, P=2.5e-5 two-sided Pearson correlation test, Fig. S31, S32).

**Fig. 5.**
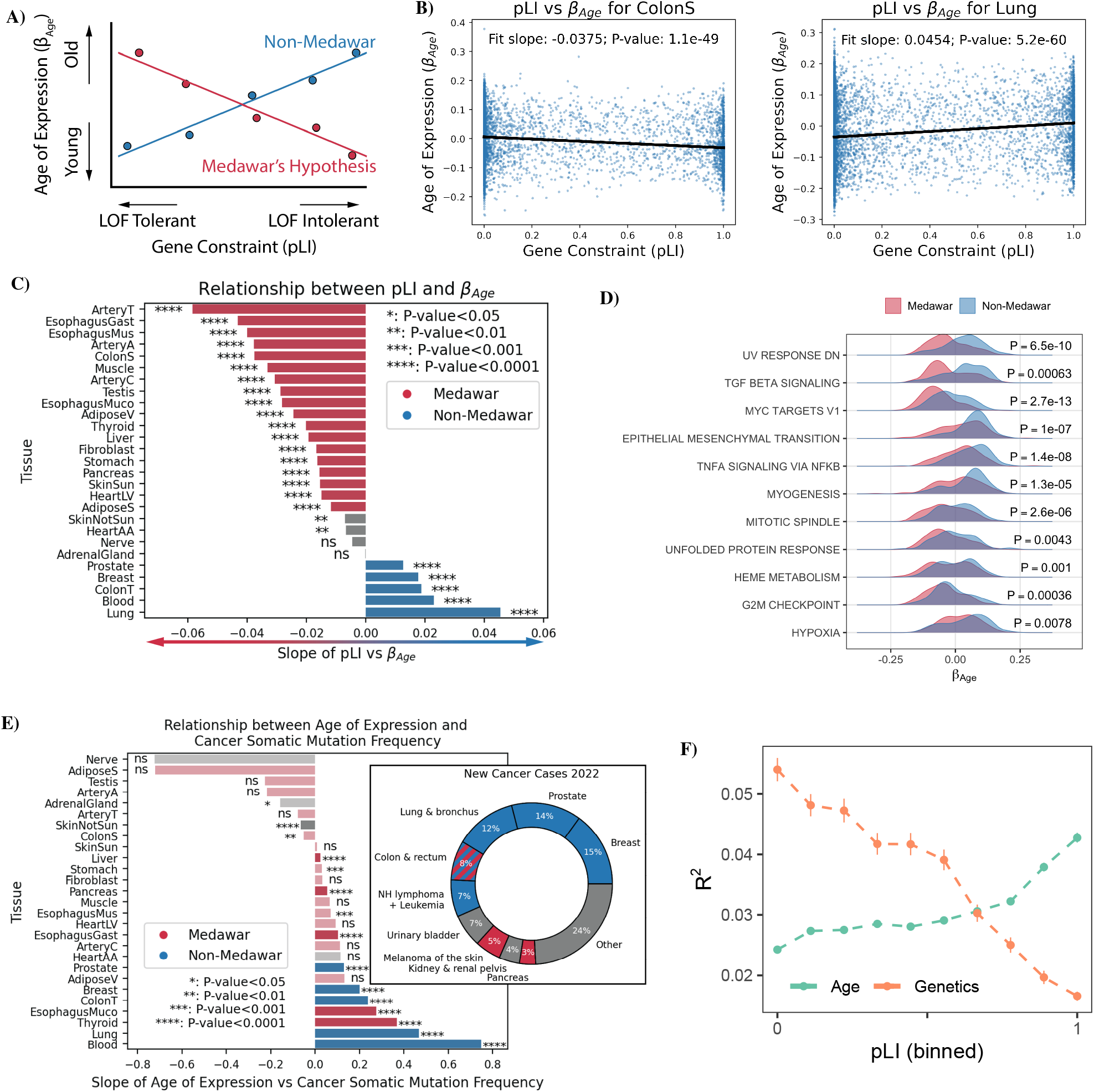
Tissue-specific evolutionary signatures of aging. **A)** The expected relationship across genes between the per-gene age-associated slope of gene expression (*β*_age_) and a genes level of constraint (measured by probability loss of function intolerance - pLI). Medawar’s hypothesis predicts a negative relationship (shown in red) between the time of expression and the level of constraint. The opposite trend (non-Medawar) is shown in blue. **B)** *β*_age_ across genes plotted as a function of pLI for a tissue exhibiting a *Medawarian* signature, and a *non-Medawarian* signature. **C)** The slope of the relationship between *β*_age_ and constraint across all tissues. Significance determined using a linear model two-sided t-test **D)** Hallmark pathways in which the *β*_age_ was significantly different between *Medawarian* and *non-Medawarian* tissues (two-sided t-test). **E)** Bar chart shows per-tissue relationship between *β*_age_ and frequency of somatic mutations in tumor samples for a particular gene and cancer type (left). Colored as in 5C with alpha indicating significance. Doughnut chart shows proportion of new cases of cancer in US in 2022 by cancer type from (23) (right)). Crosshatch indicates that while colon transverse was identified as a non-Medawar tissue, colon sigmoid was not. **F)** Gene expression variance explained by genetics and age as a function of binned gene constraint averaged across all tissues.

To explore why these five tissues might exhibit distinctive evolutionary signatures of aging we compared the distribution of significant *β*_age_ parameters between *Medawarian* and *non-Medawarian* tissues among different *hallmark pathways* (22). We found 11 signatures exhibiting significantly increased *β*_age_ (FDR<0.01 two-sided t-test) compared to *non-Medawarian* tissues (Fig. 5D, S33) including DNA-damage, TGF-*β* signalling, MYC targets, and epithelial-to-mesenchymal transition pathways most prominently. All of these signatures are broadly correlated with cellular proliferation, differentiation, and cancer. Indeed, these five *non-Medawarian* tissues are also the top five most commonly diagnosed sites for cancer in 2022 (23) (Fig. 5E). To directly investigate cancer signatures in these tissues we quantified the per-gene likelihood of having somatic mutations in tumors using the COSMIC cancer browser (24). GTEx tissues were matched to most representative cancer types for comparisons (e.g. Breast Cancer *→* Breast Mammary Tissue, Table S6). We found that the per-gene age of expression (*β*_*age*_) was significantly correlated with mutation frequency (i.e. mutational burden) across several tissues (Fig. 5E, S34) with the 5 *non-Medawarian* tissues exhibiting some of the strongest signatures (P<10^−4^ linear model two-sided t-test). These results highlight that gene expression patterns in tissues and celltypes that proliferate throughout the course of an individuals life may be subjected to distinct evolutionary pressures with important implications for the cancer susceptibility of these tissues.

We also explored the relationship between gene expression heritability and constraint. Across all tissues *h*^2^ was significantly negatively correlated with pLI (P-value < 10^−3^ linear model two-sided t-test, Fig. S35, S36, S37). While this trend was consistent across tissues, intriguingly it was strongest in heart tissues. Thus, on average genes in which the variation in expression levels is heritable tend to be under significantly less functional constraint (Fig. 5F). In contrast however, on average 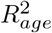 increases as function of pLI (Fig. 5F), highlighting the increased constraint of many of the genes that exhibit age-associated changes in gene expression. These highly conserved genes (e.g. the aforementioned mitochondrial and translation factors) are thus potentially of critical importance to disease. Together, these results highlight the stark contrast in the types of genes with heritable expression patterns (reduced constraint) compared to those with age-associated gene expression patterns (increased constraint.

## Discussion

Studying age-associated changes in gene expression provides critical insights into the underlying biological processes of aging. Here, we set out to quantify the relative contributions of aging and genetics to gene expression phenotypes across different human tissues. Our study finds that the predictive power of eQTLs is significantly impacted by age across several different tissues and that this effect is more pronounced in older individuals. These results extend upon previous work examining blood tissue (4) and highlight the varied impact of aging on eQTLs among different tissues. We show that this result is likely to be in part due to an increase in the inter-individual heterogeneity of gene expression patterns among individuals in some contexts, potentially as a result of the increased impact of the environment. It is also of note that increased inter-individual heterogeneity in both younger and older individuals was associated with reduced predictive power of eQTLs. However, our study is limited in it’s primary focus on bulk-tissue transcriptomic data. Early evidence from single cell studies already suggests that differences in gene expression heterogeneity vary among cell types of tissues as a function of age (6, 7, 25, 26). While these studies lack sufficient individual sample sizes and genetic diversity for the statistical approaches used herein, it is possible that in the future the availability of larger datasets will facilitate studying these phenomena at the single-cell level. The extensive tissue heterogeneity we observe suggests that patterns of aging will exhibit substantial cell-type specificity.

We also present a novel approach to jointly model the impact of genetics and aging on gene expression variance to parse out the individual contributions of each of these factors. The increased complexity of our model has little impact on its accuracy with our expression heritability estimates strongly correlated with previous heritability measures across all tissues (mean Pearson’s r=0.89, Fig. S18). Using this model we show that age exhibits exceptionally varied affects on different tissues, and indeed, in several tissues age contributes more to gene expression variance on average than genetics. These results also highlight a widespread coordinated signature of age-associated decline in mitochondrial and translation factors. Dysregulation in mitochondrial function and ribosome biogenesis have been documented as key players in aging, (27, 28), however our results highlight the tissuespecificity of these trends. Our model also allows us to quantify the tissue-specific evolutionary context of age-associated gene expression changes. We corroborate the inverse relationship between age-at-expression and constraint, as predicted by Medawar’s hypothesis and recently documented by others (8, 9, 19) across the vast majority of tissues. However, we also surprisingly identify five tissues which exhibit the opposite pattern and show that age-associated signatures of increased proliferation and cancer are enriched in these tissues. These results highlight the distinct evolutionary forces that act on late-acting genes expressed in highly proliferative celltypes. Future work extending these analyses to the single-cell level will provide further insights into the cell-type-specific age-associated patterns of constraint, and its relevance to cancer.

Overall this work has several important implications. Our results shed light on recent work on the prediction accuracy of polygenic risk scores (PRS) (29) which found that numerous factors, including age, sex, and socioeconomic status can profoundly impact the prediction accuracy of such scores even in individuals with the same genetic ancestry. Our results highlight that genetics exhibit varied predictive power in several different tissues as a function of age, potentially playing a role in differential PRS accuracy between young and old individuals. This also has important implications for disease association and prediction approaches that leverage expression quantitative trait loci to prioritize variants, including colocalization methods (30), transcriptome-wide association studies (14, 31), and Mendelian randomization (32, 33). If a significant proportion of eQTLs exhibit age-associated biases in their effect size in a tissue of interest, then these approaches may be less powerful when applied to diseases for which age is a primary risk factor such as heart disease, Alzheimer’s dementias, cancers, and diabetes. Furthermore our results highlight that genes with eQTLs tend to be subject to less evolutionary constraint, and thus potentially less biologically important, in contrast to genes with age-associated gene expression patterns which exhibit increased constraint.

The critical role of aging as a risk factor for many common human diseases underscores the importance of understanding its impact on cellular systems at the molecular level. Together our analyses provide novel insights into tissue-specific patterns of aging and the relative impact of genetics and aging on gene expression. We anticipate that future studies across tissues and cells of gene expression, chromatin structure, and epigenetics will further elucidate how both programmed and stochastic processes of aging drive human disease.

## Supplementary Note 1: Methods

### Data collection age groupings

We downloaded gene expression data for multiple individuals and tissues from GTEx V8 (10), which were previously aligned and processed against the hg19 human genome. Tissues were included in the analysis if they had >100 individuals in both the age ≥55 and <55 cohorts described below (Fig. S2). For a given tissue, genes were included if they had >0.1 TPM in ≥ 20% of samples and ≥ 6 reads in ≥ 20% of samples, following GTEx’s eQTL analysis pipeline. To compare gene expression heritability across individuals of different ages, for some analyses we split the GTEx data for each tissue into two age groups, “young” and “old,” based on the median age of individuals in the full dataset, which was 55 (Fig. S1). Within each tissue dataset, we then equalized the number of individuals in the young and old groups by randomly downsampling the larger group, to ensure that our models were equally powered for the two age groups.

### PEER factor analysis

We analyzed existing precomputed PEER factors available from GTEx to check for correlations between these hidden covariates and age. In particular, we fit a linear regression between age and each hidden covariate and identified significant age correlations using an F-statistic (Fig. S3). Because some of the covariates were correlated with age, we generated new age-independent hidden covariates of gene expression to remove batch and other confounding effects on gene expression while retaining age related variation. In particular, we first removed age contributions to gene expression by regressing gene expression on age and then ran PEER on the age-independent residual gene expression to generate 15 age-independent hidden PEER factors.

### Quantifying the effect of eQTLs on gene expression in different age groups

Using the binary age groups defined above, we assessed the relative significance of eQTLs in old and young individuals by carrying out separate assessment of eQTLs identified by GTEx. We report the number of genes included in analysis for each tissue (Table S7). For each gene in each tissue and each age group, we regressed the GTEx pre-normalized expression levels on the genotype of the lead SNP (identified by GTEx, MAF>0.01) using 5 PCs, 15 PEER factors, sex, PCR protocol and sequencing platform as covariates, following the GTEx best practices. We confirmed our results using both our recomputed PEER factors as well as the PEER factors provided by GTEx (Fig. S4). To test for significant differences in genetic associations with gene expression between the old and young age groups, we compared the p-value distributions between these groups for all genes and all SNPs in a given tissue using Welch’s t-test. To investigate the validity of the age cutoff used for these binary age groups, we replicated the eQTL analysis using two additional age cutoffs of 45 and 65 years old. We observed the same trends in both cases; however, statistical power decreased due to smaller sample sizes in the resulting age bins, leading to a non-significant result for age cutoff 45 (Fig. S38).

### Jensen-Shannon Divergence as a distance metric between transcriptome profiles

To quantify differences in gene expression between individuals, we computed the pairwise distance for all pairs of individuals in an age group using the square root of Jensen-Shannon Divergence (JSD) distance metric, which measures the similarity of two probability distributions. Here we applied JSD between pairs of individuals’ transcriptome vectors containing the gene expression values for each gene, which we converted to a distribution by normalizing by the sum of the entries in the vector. For two individuals’ transcriptome distributions, the JSD can be calculated as:

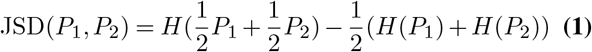

where *P*_*i*_ is the distribution for individual *i* and H is the Shannon entropy function:

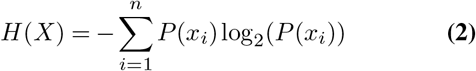

JSD is known to be a robust metric that is less sensitive to noise when calculating distance compared to traditional metrics such as Euclidean distance and correlation. It has been shown that JSD metrics and other approaches yield similar results but that JSD is more robust to outliers (12). The square root of the raw JSD value follows the triangle inequality, enabling us to treat it as a distance metric.

### Slope of JSD distance versus age

In addition to comparing JSD between the two age groups defined above, “young” and “old”, we also binned all GTEx individuals into 6 age groups, from 20 to 80 years old with an increment of 10 years. We then computed pairwise distance and average age for each pair of individuals within each bin using the square root of JSD as the distance metric. We applied a linear regression model of JSD versus age to obtain slopes, confidence intervals, and p-values.

### Cell-type specific analysis

To analyze whether cell type composition affects age-associated expression changes, we utilized the tool CIBERSORTx (16) to estimate cell type composition and individual cell type expression levels in GTEx whole blood. Cell type composition estimates were computed using CIBERSORTx regular mode. Individual cell type expression level estimates were computed using CIBER-SORTx high resolution mode. We then repeated our JSD and eQTL analyses on each cell type independently (see JSD and eQTL sections for details). In addition, to analyze tissuespecific differences in cell type composition, we referred to a previous study (34) that computed cell type composition for different GTEx tissues using CIBERSORTx. We applied the JSD metric to each tissue, using the cell type composition vector as the distribution. Additionally, we applied the Breusch–Pagan test to compute heteroskedasicity coefficients and p-values with respect to age, after inverse logit transformation to give an approximately Gaussian distribution (Fig S42) (see section on heteroskedastic gene expression).

### Heteroskedastic gene expression

We used the Breusch–Pagan test to call heteroskedastic gene expression with age. For each gene and tissue, we computed gene expression residuals by regressing out age-correlated PEER factors, other GTEx covariates, and age. To test for age-related heteroskedasticity, we squared these residuals and divided by the mean, regressed them against age, and looked at the age effect size (*β*_*het*_). We called significantly heteroskedastic genes using a two-sided t-test with the null hypothesis that the *β*_*het*_ is zero. The Benjamini-Hochberg procedure was used to control for false positives. To determine which tissues have more genes with increasing gene expression heterogeneity with age, we compare the number of genes with positive heteroskedasticity (*β*_*het*_>0 and *FDR*<0.2) to the total of all heteroskedastic genes (*FDR*<0.2). We compare this metric to the per-tissue 2-bin JSD (Fig. S39) and 6-bin JSD slope (Fig. S13).

### Multi-SNP gene expression prediction

We used a multiSNP gene expression prediction model based on PrediXcan (14) to corroborate our findings from the eQTL and JSD analyses on the two age groups, “young” and “old”. For each gene in each tissue, we trained a multi-SNP model separately within each age group to predict individual-level gene expression.

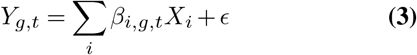

Where *β*_*i,g,t*_ is the coefficient or effect size for SNP *X*_*i*_ in gene *g* and tissue *t* and *c* includes all other noise and environmental effects. The regularized linear model for each gene considers dosages of all common SNPs within 1 megabase of the gene’s TSS as input, where common SNPs are defined as MAF > 0.05 and Hardy-Weinberg equilibrium P > 0.05. We removed covariate effects on gene expression prior to model training by regressing out both GTEx covariates and age-independent PEER factors (described above). Coefficients were fit using an elastic net model which solves the problem ((35)):

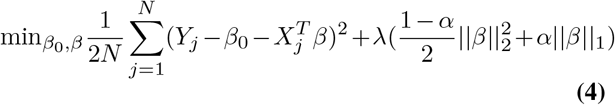

The minimization problem contains both the error of our model predictions 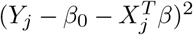 and a regularization term 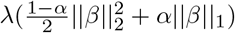 to prevent model overfitting. The elastic net regularization term incorporates both L1 (| |*β*| | _1_)) and 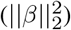 penalties. Following PrediXcan, we weighted the L1 and L2 penalties equally using *α* = 0.5 (14). For each model, the regularization parameter *λ* was chosen via 10-fold cross validation. The elastic net models were fit using Python’s glmnet package and *R*^2^ was evaluated using scikit-learn. From the trained models for each gene, we evaluated training set genetic *R*^2^ (or *h*^2^) for the two age groups and subtracted 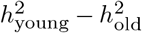 to get the difference in gene expression heritability between the groups. We compared this average difference in heritability to the mean JSD_*old*_ − JSD_*young*_ and log(*P*_*old*_) − log(*P*_*young*_) using P-values from the eQTL analyses across genes.

### Joint model for expression prediction using SNPs and age

To uncover linear relationships between gene expression and both age and genetics, we built a set of gene expression prediction models using both common SNPs and standardized age as input. An individual’s gene expression level *Y* for a gene *g* and tissue *t* is modeled as:

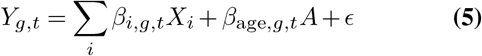

Where *A* is the normalized age of an individual. Coefficients were fit using elastic net regularization, as above, which sets coefficients for non-informative predictors to zero. The sign of the fitted age coefficient (*β*_*age,g,t*_), when nonzero, reflects whether the gene in that tissue is expressed more in young (negative coefficient) or old (positive coefficient) individuals. We also evaluated the training set *R*^2^ using the fit model coefficients separately for genetics (across all SNPs in the model) and age:

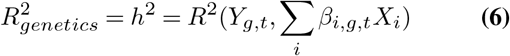

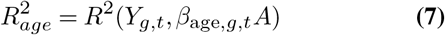

We also tested whether the age-related gene expression relationship was sex-specific by rerunning the joint model with an additional age-sex interaction term as follows:

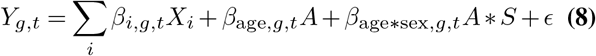

Where *β*_age*sex,*g,t*_ is the additional model weight for the agesex interaction term and *S* is the binary sex of the GTEx individual. The *R*^2^ of age, genetics, and the age-sex interaction term are evaluated as before by determining the variance explained by each term in the model. We compared the 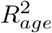 between the models including or excluding the age-sex interaction term (Fig. S24). We also compared the tissueaveraged variance explained by age and the age-sex interaction term. Finally, to check the consistency of tissue-specific gene expression heritability estimates from our model and the original PrediXcan model trained on GTEx data, we evaluate Pearson’s r between our heritability estimates and those of PrediXcan (Fig. S18), using heritability estimates from the original PrediXcan model available in PredictDB.

### Tissue specificity of age and genetic associations

We evaluated the variability of age and genetic associations across tissues using a measure of tissue specificity for age and genetic *R*^2^ (36). We measured the tissue-specificity of a gene *g*’s variance explained 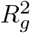 using the following metric:

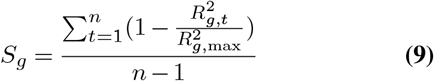

Where *n* is the total number of tissues, 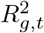 is the variance explained by either age or genetics for the gene *g* in tissue *t* and 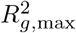 is the maximum variance explained for *g* over all tissues. This metric can be thought of as the average reduction in variance explained relative to the maximum variance explained across tissues for a given gene. The metric ranges from 0 to 1, with 0 representing ubiquitously high genetic or age *R*^2^ and 1 representing only one tissue with nonzero genetic or age *R*^2^ for a given gene. We calculate *S*_*g*_ separately for 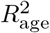 and 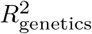 across all genes.

### Functional constraint analysis

We quantified gene constraint using the probability of loss of function intolerance (pLI) from gnomAD 2.1.1 (20). We analyzed the relationships between pLI vs *β*_age_ and pLI vs heritability across genes. For these analyses, genes were only included if age or genetics were predictive of gene expression (*R*^2^ > 0) for that gene. For genes with *R*^2^ > 0, we used linear regression to determine the direction of the relationship between pLI and *β*_age_ or heritability for each tissue. The F-statistic was used to determine whether pLI was significantly related to these two model outputs. For pLI vs *β*_age_, a significant negative slope was considered a *Medawarian* trend (consistent with Medawar’s hypothesis) and a significant positive slope a *non-Medawarian* trend. To test whether the non-Medawarian trends were driven by genes with higher expression, we excluded genes in the top quartile of median gene expression and repeated the analysis between pLI and *β*_age_ (Fig. S40). We also analyzed the evolutionary constraint metric dN/dS (21) and its tissue-specific relationship with *β*_age_ by determining the slope and significance of the linear regression, as above.

### Cancer Somatic Mutation Frequency

We quantified the per-gene and per-tissue cancer somatic mutation frequency using data from the COSMIC cancer browser (24). For each tissue, we selected the closest cancer type as noted in Table S5 and downloaded the number of mutated samples (tumor samples with at least one somatic mutation within the gene) and the total number of samples for all genes. We computed the cancer somatic mutation frequency by dividing the number of mutated samples by the total number of samples. For each tissue, we plotted the gene’s *β*_age_ vs its cancer somatic mutation frequency for all genes with >200 tumor samples. We report the slope and significance of the relationship between *β*_age_ and cancer somatic mutation frequency for each tissue. To determine whether age-dependent gene expression heteroskedasticity is related to a gene’s involvement in cancer (Fig. S41), we also plotted each gene’s heteroskedasticity effect size vs the cancer somatic mutation frequency for all genes with >200 tumor samples and moderately significant heteroskedasticity (FDR<0.2). Tissues with ≤5 genes meeting these criteria are not plotted.

### *Non-Medawarian* tissue analysis

To explore the *nonMedawarian* trend in some tissues, we assessed the distribution of *β*_age_ across *Medawarian* and *non-Medawarian* tissues for genes within each of the 50 MSigDB hallmark pathways (22). Significant differences between the distributions were called using a t-test, and p-values were adjusted for multiple hypothesis testing using a Benjamini-Hochberg correction.

## Data availability

All data are available on GitHub https://github.com/sudmantlab/gene_expression_aging and full results for joint age and genetic model can be found on Zenodo https://doi.org/10.5281/zenodo.6555453.

## Code availability

All analyses were performed in R version 4.0.2 and Python 3.6. All code is available online at https://github.com/sudmantlab/gene_expression_aging and archived at https://doi.org/10.5281/zenodo.6555500.

## ACKNOWLEDGEMENTS

This work was supported by the National Institute of General Medical Sciences grant R35GM142916 to P.H.S. and the National Human Genome Research Institute grant R00HG009677 to N.M.I.

## AUTHOR CONTRIBUTIONS

RY, RC, JMV, HS, PS, and PHS performed all analysis. RY, RC, NMI, and PHS wrote the manuscript. PHS and NMI supervised the project. PHS conceived of the project.

## Supplementary Figures

**Fig S 1.**
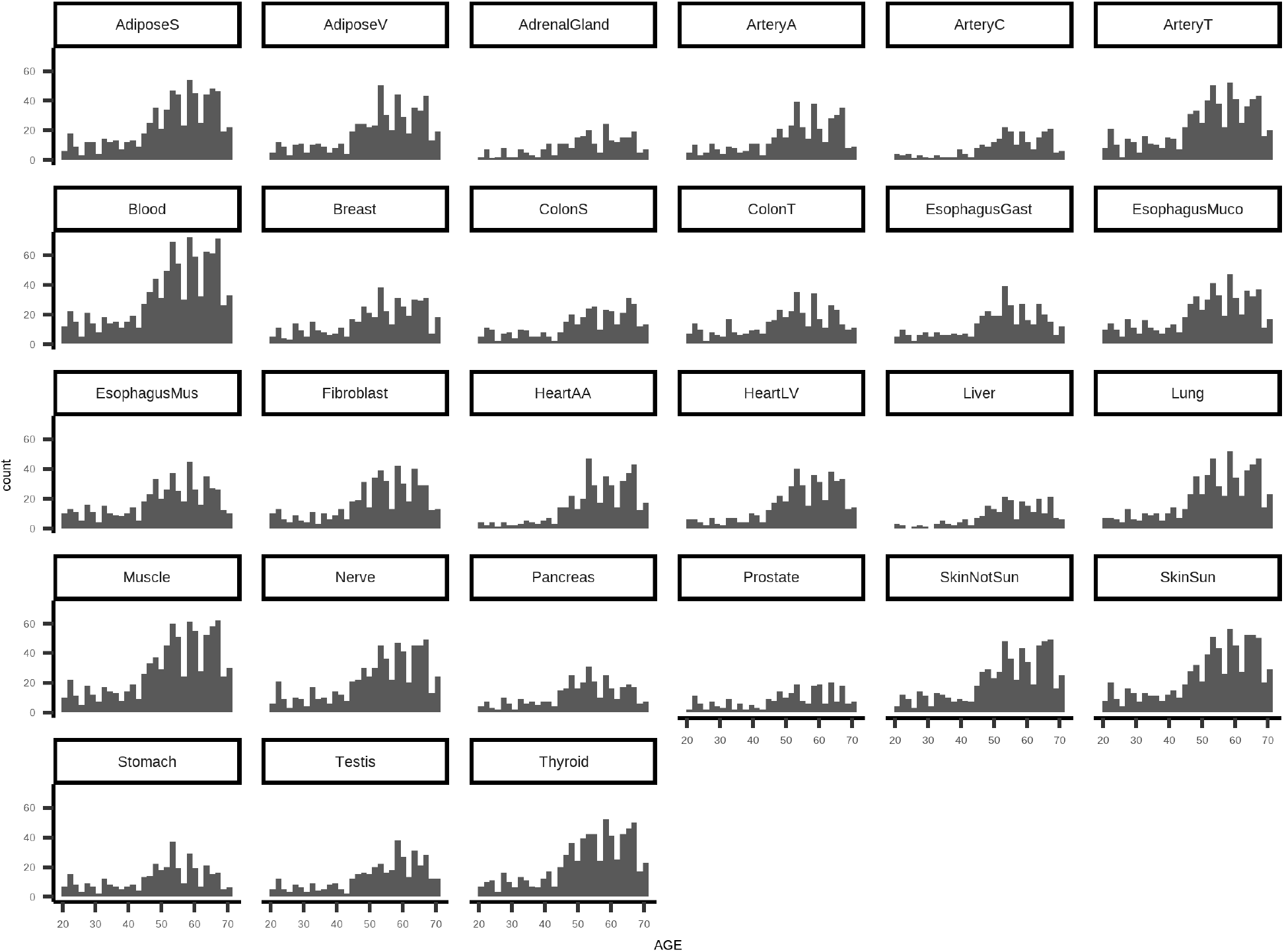
Histogram representing age distribution of each GTEx tissue, x-axis representing age of each sample and y-axis representing counts in bin.

**Fig S 2.**
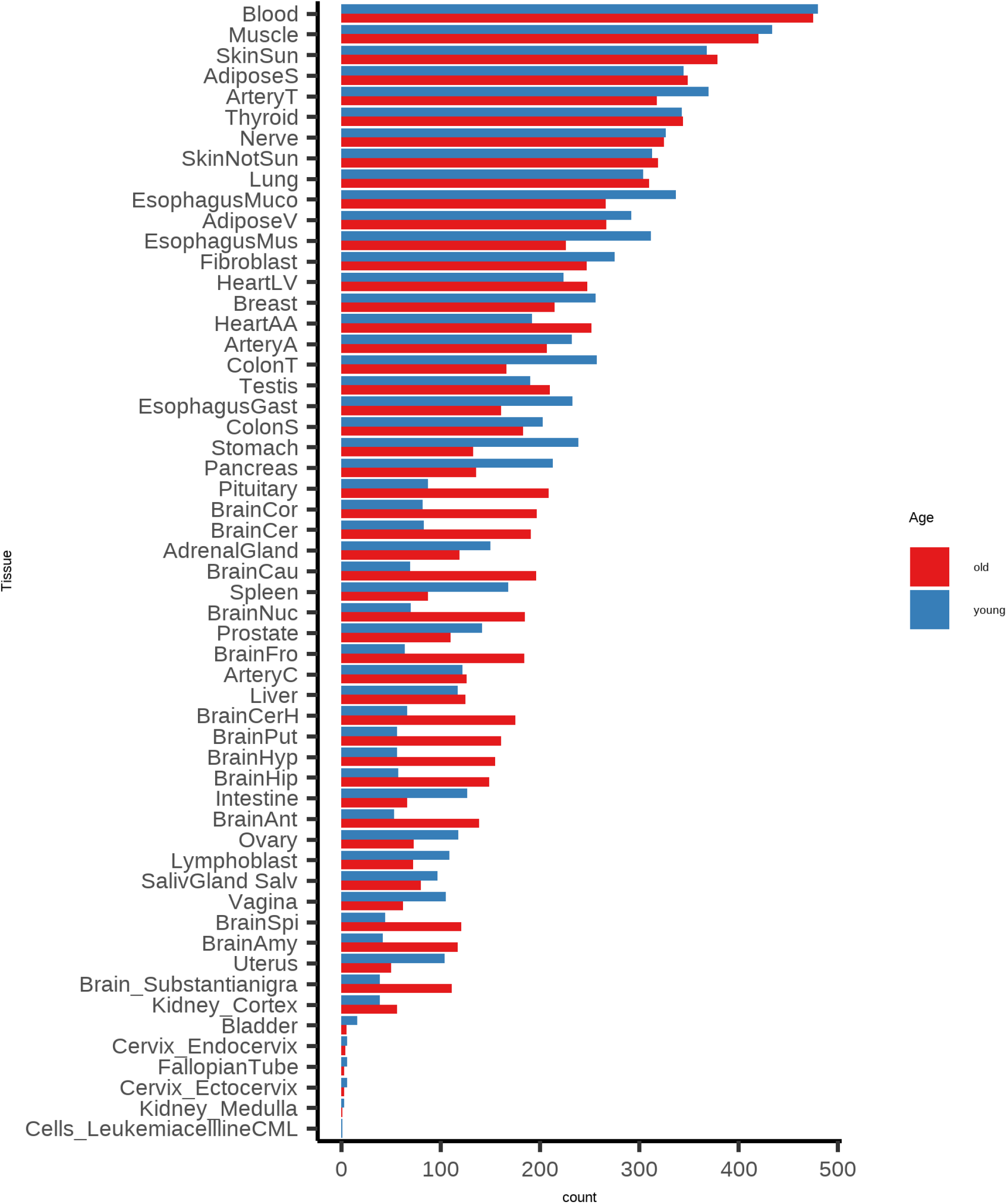
Number for individuals above and below the median age of 55 by tissue for all 47 GTEx tissues.

**Fig S 3.**
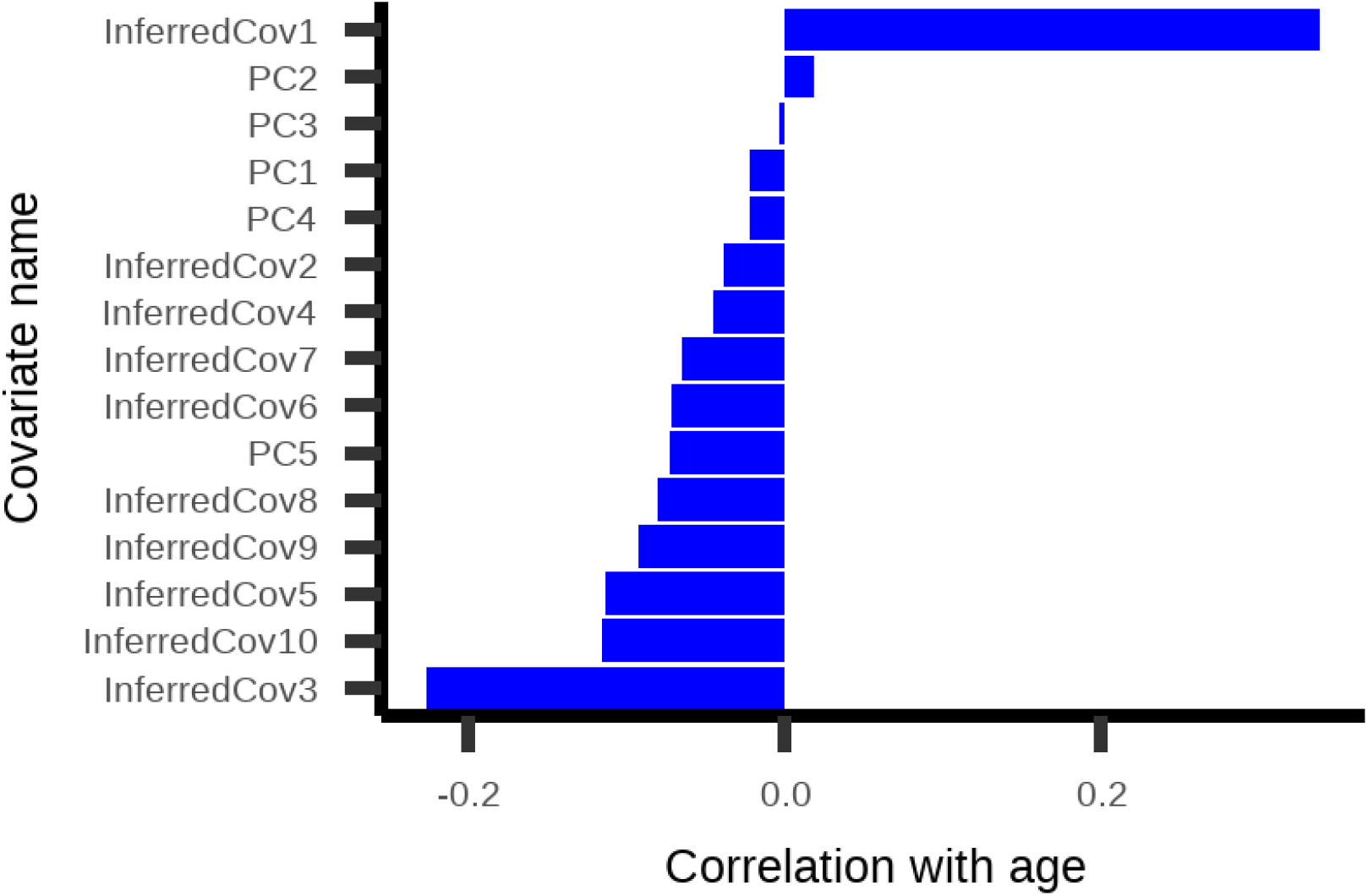
Pearson’s r between GTEx covariates with age. PEER factor covariates labeled as “InferredCov” and genetic principle components labeled as “PC”

**Fig S 4.**
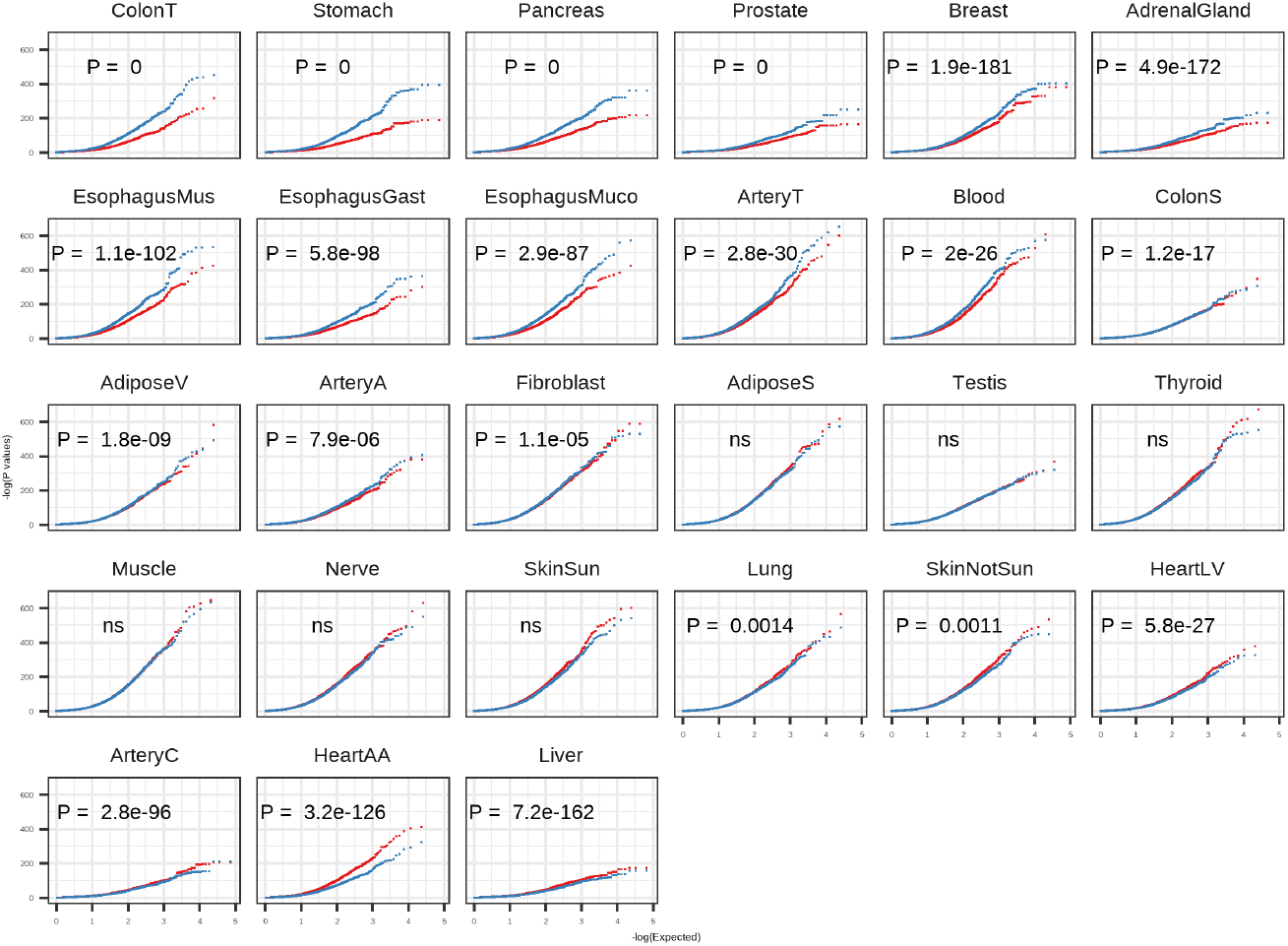
QQ plot of eQTL p-values for old (red) and young (blue) cohorts across 27 tissues using GTEx PEER factors. Significant differences between p-value distributions annotated for each tissue. P-values are obtained from two-sided Welch’s t test.

**Fig S 5.**
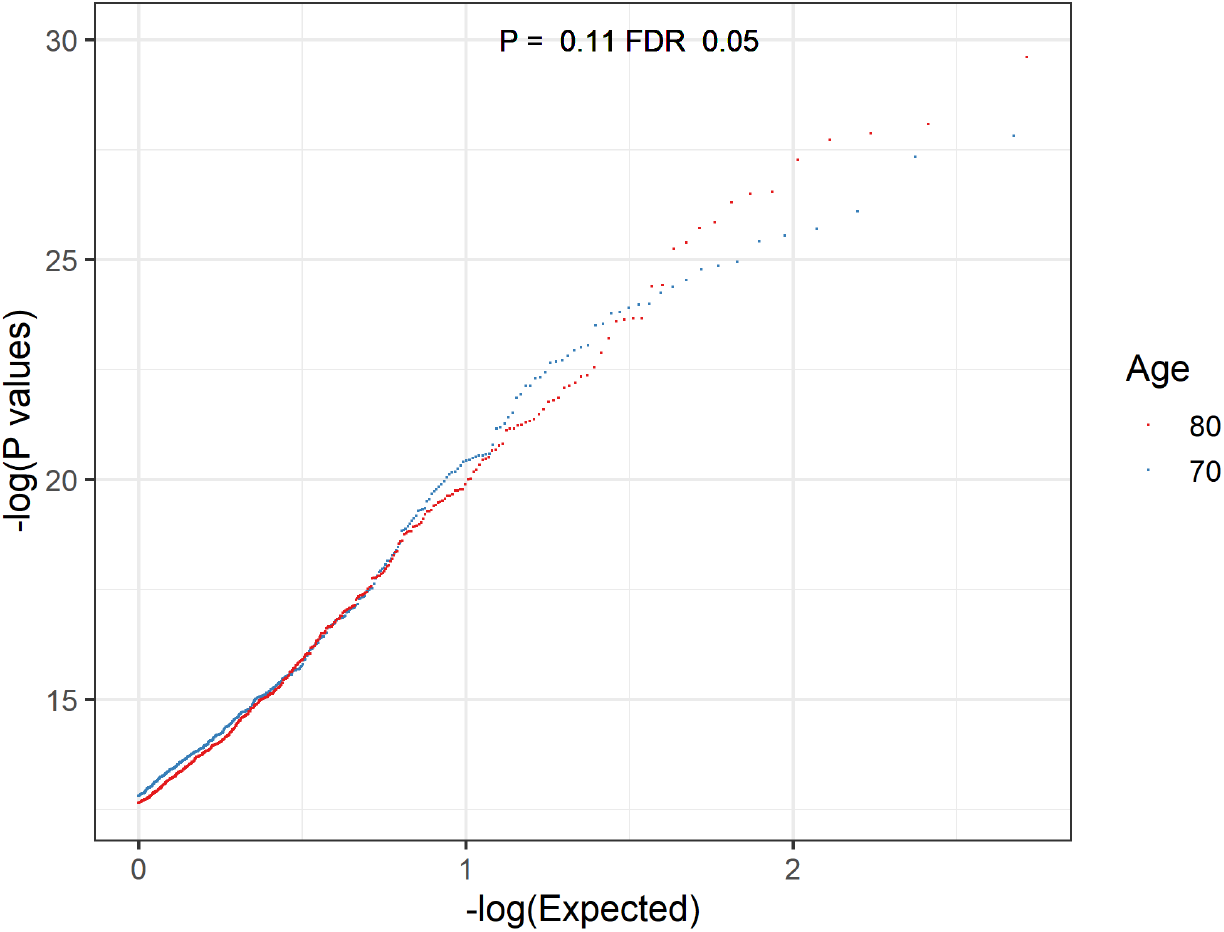
QQ plot of eQTL p-values from individuals of age 70 and 80 from PIVUS cohort using genes under FDR cutoff of 0.05. P-values for distribution difference are obtained from two-sided Welch’s t test.

**Fig S 6.**
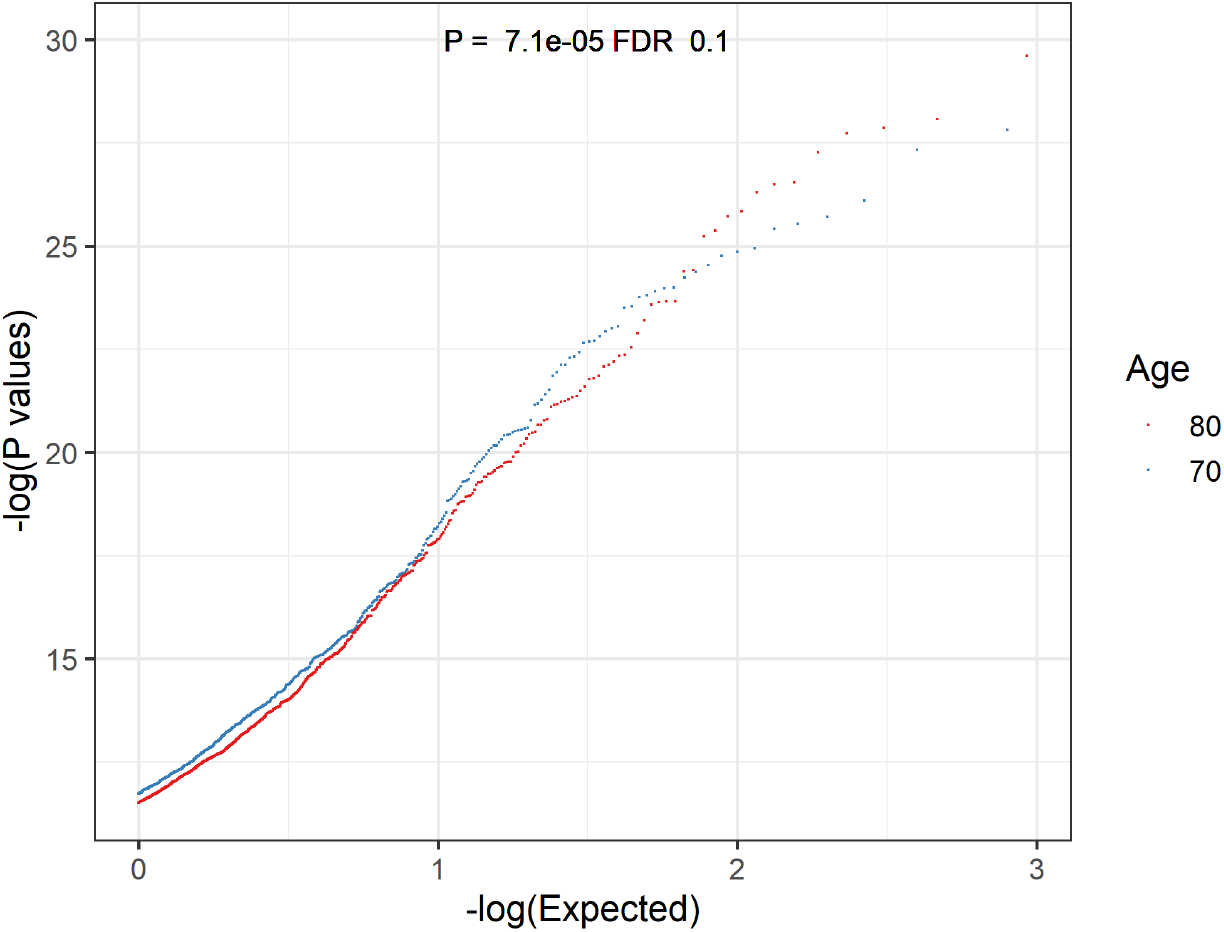
QQ plot of eQTL p-values from individuals of age 70 and 80 from PIVUS cohort using genes under FDR cutoff of 0.1. P-values for distribution difference are obtained from two-sided Welch’s t test.

**Fig S 7.**
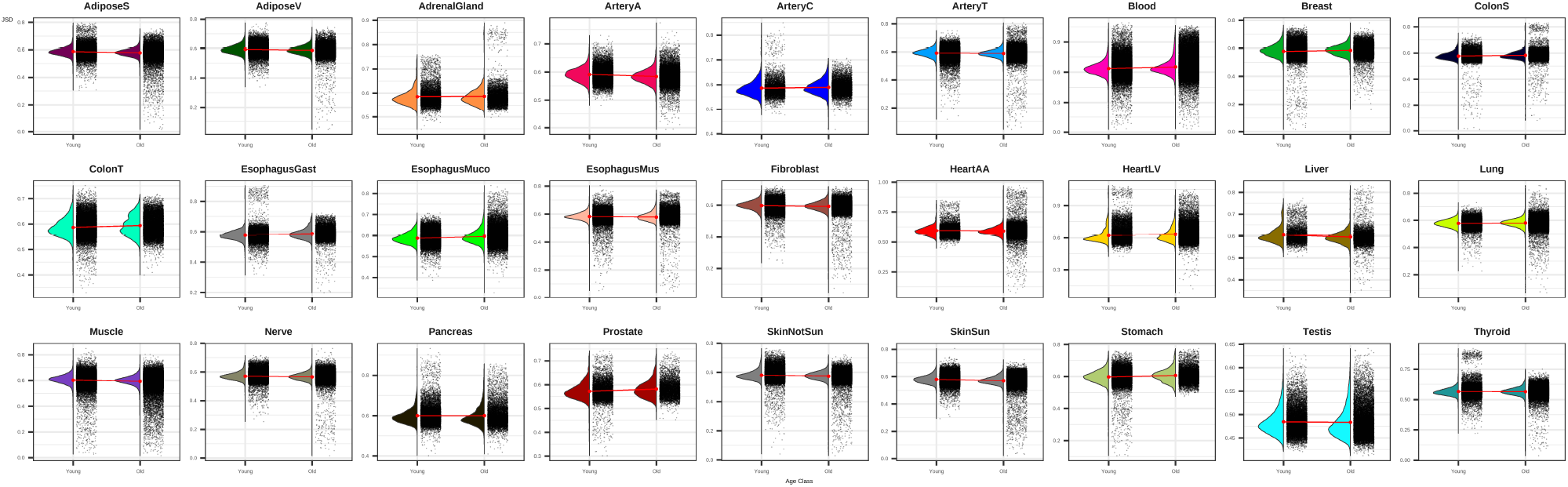
The distributions of JSD distances for all tissues in old and young bins.

**Fig S 8.**
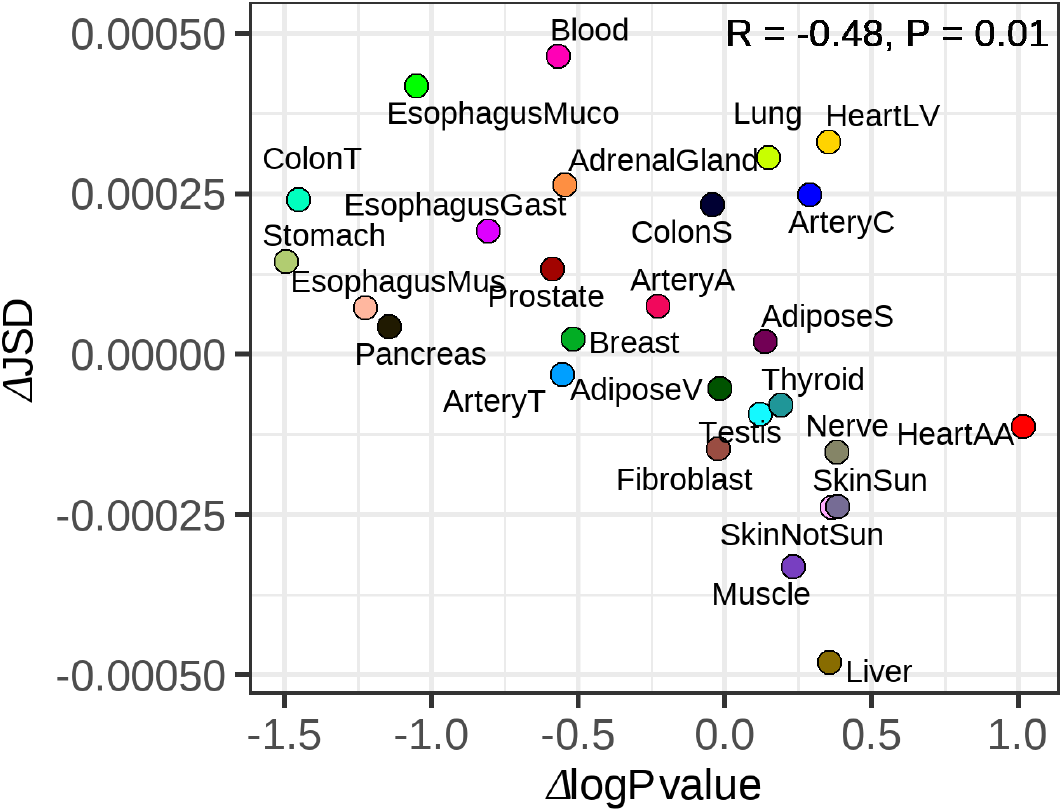
Scatter plot showing the correlation between the difference in average JSD between young and old individuals and difference in eQTL log(p-value) between young and old. R and p-value are obtained from pearson correlation test.

**Fig S 9.**
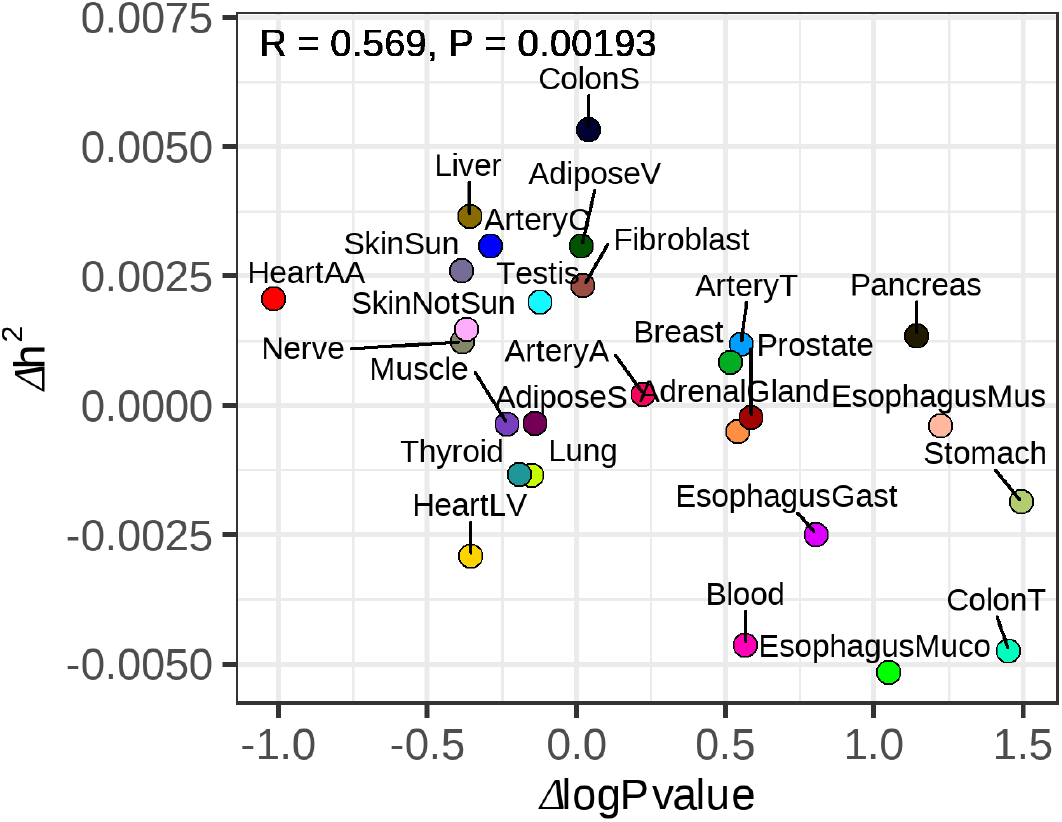
High correspondence of age-related changes in gene expression associations for single SNP and multi-SNP models. Scatter plot showing the correlation between the difference in heritability estimated by the multi-SNP linear model (PrediXcan) in young and old individuals and difference in eQTL log(p-value) in young and old individuals.R and p-value are obtained from pearson correlation test.

**Fig S 10.**
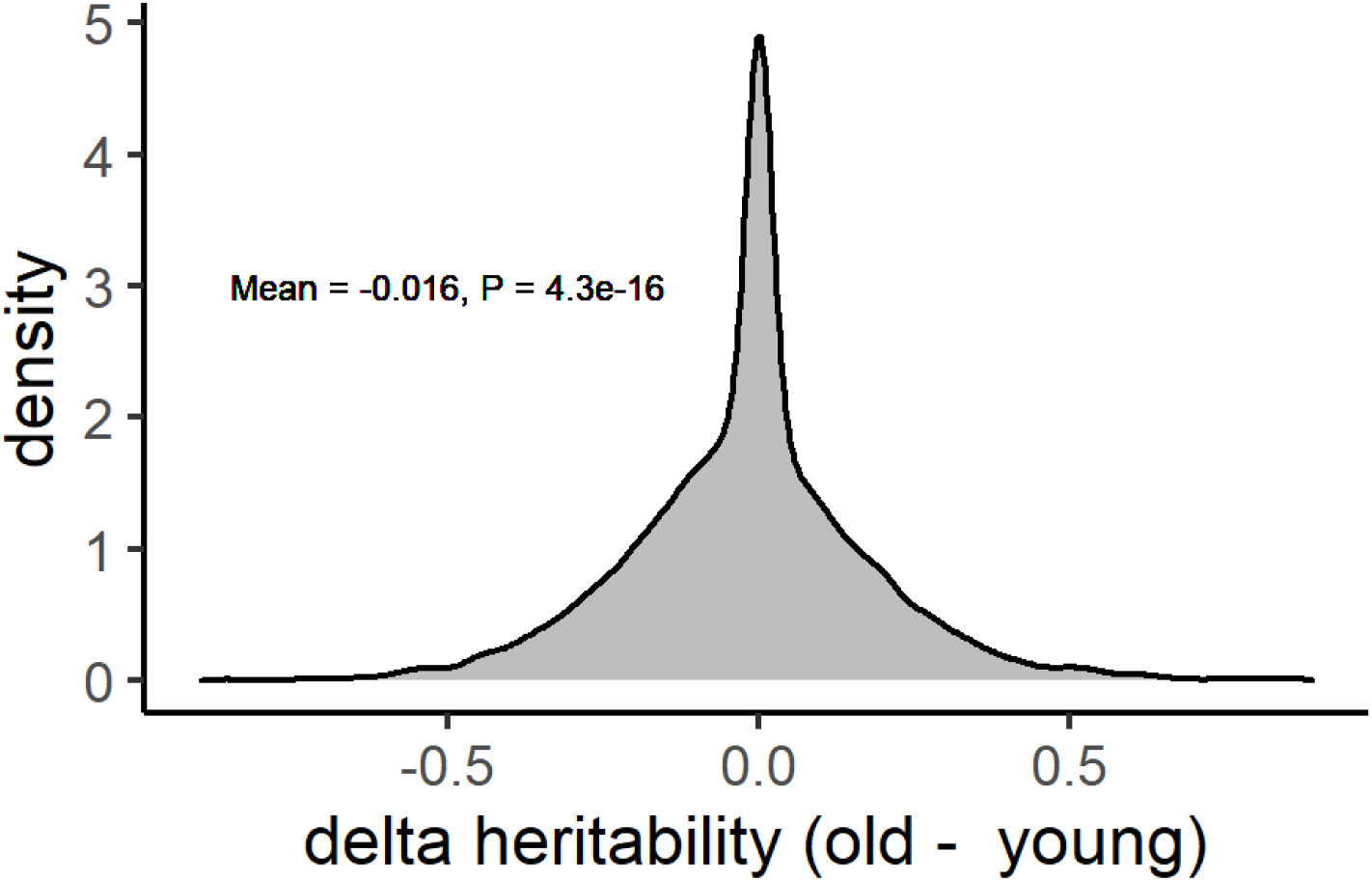
Distribution of Δ*h*^2^ (80 - 70) in PIVUS cohort. P-value is obtained from two-sided one-sample t-test.

**Fig S 11.**
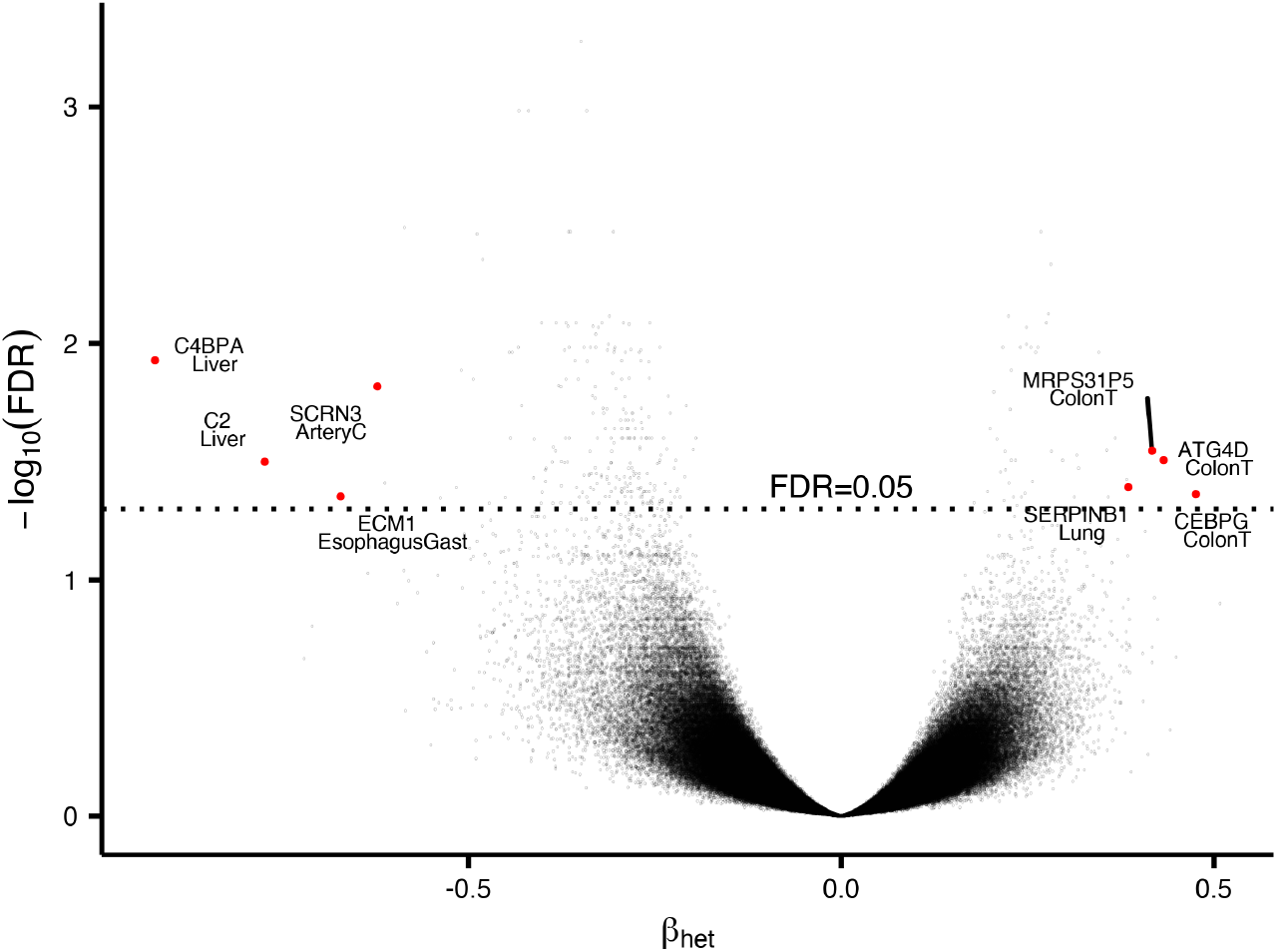
Volcano plot of the age-related heteroskedasticity significance vs effect size for each tested gene’s expression. FDR cutoffs of 0.05 is indicated. Genes with top 4 largest and smallest effect sizes with significant heteroskedasticity are labeled.

**Fig S 12.**
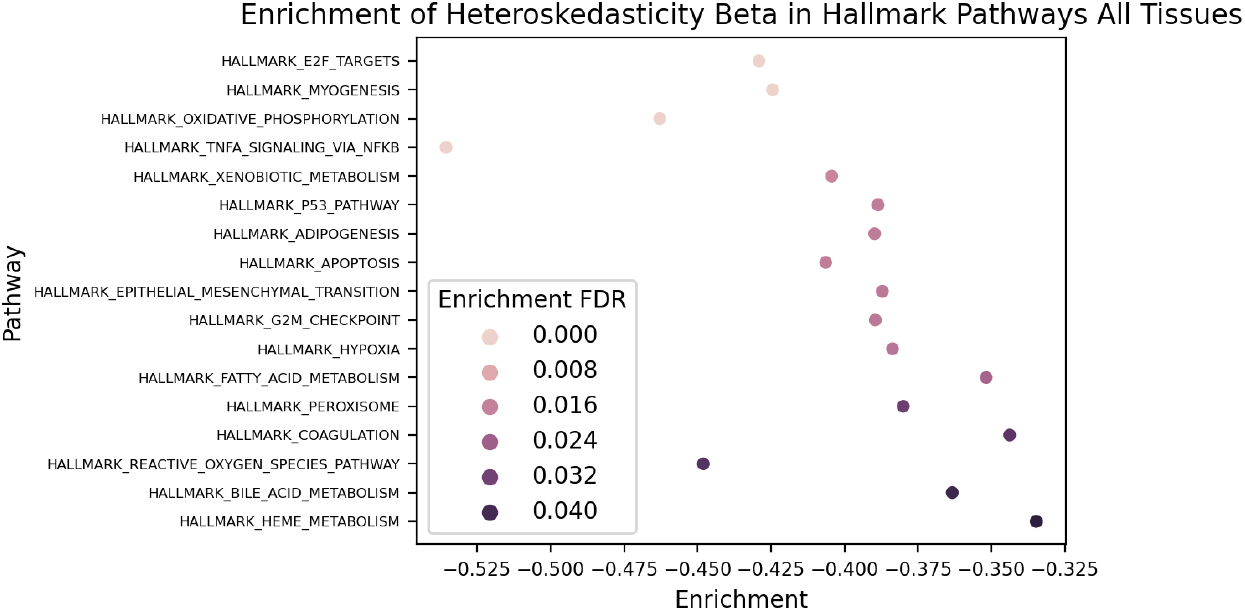
Gene set enrichment analysis of significantly heteroskedastic genes (FDR<0.05) ranked by tissue-averaged heteroskedasticity effect size using the MSigDB hallmark pathway gene sets.

**Fig S 13.**
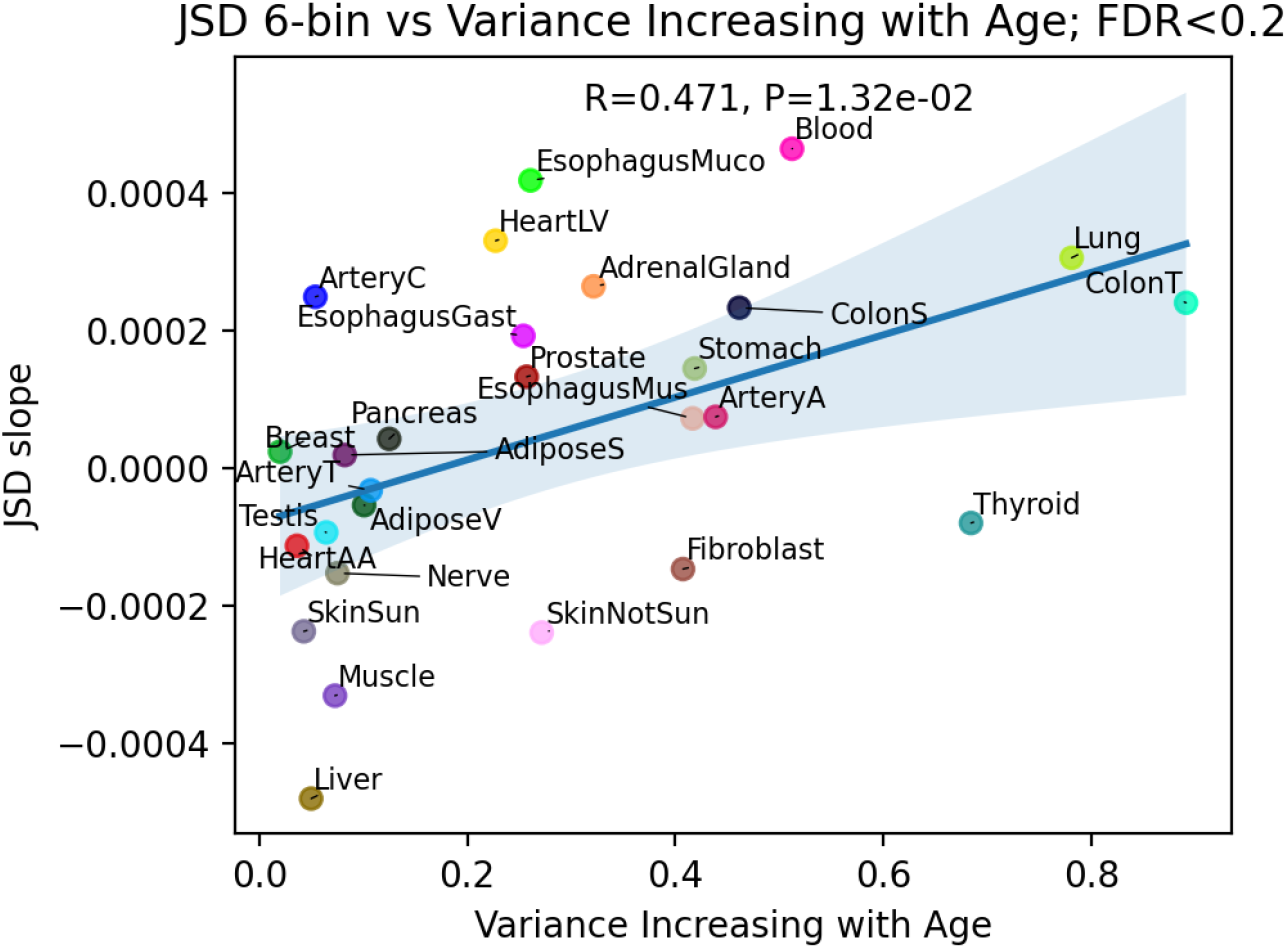
JSD age-related heterogeneity metric with 6 age bins vs proportion of heteroskedastic genes with increasing expression variance with age.

**Fig S 14.**
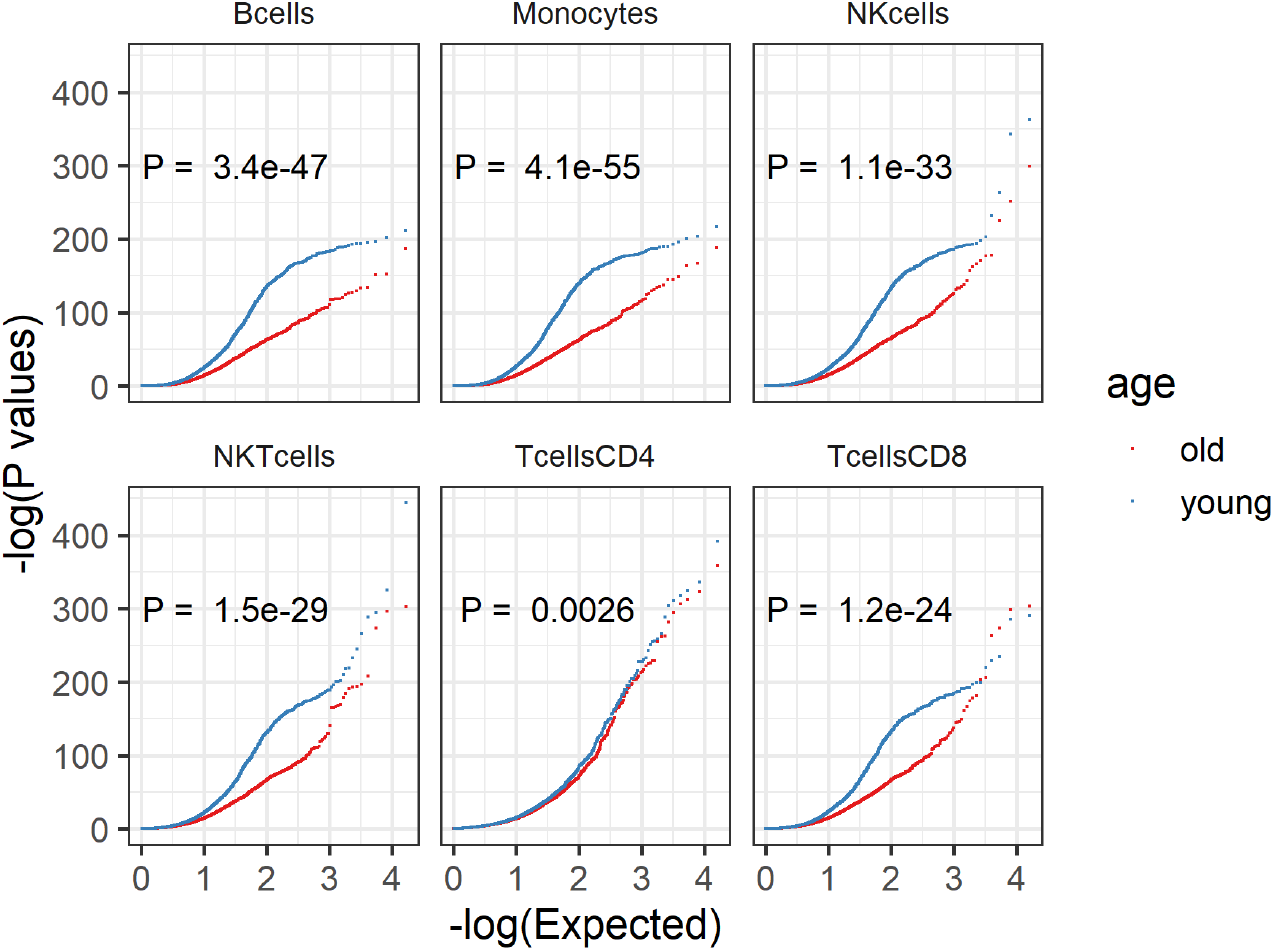
QQ plot of eQTL p-values from cell type imputed gene expression profiles of GTEx Whole Blood. Cell type expression levels are estimated from CIBERSORTx high resolution mode using all genes. P-values for distribution difference are obtained from two-sided Welch’s t test.

**Fig S 15.**
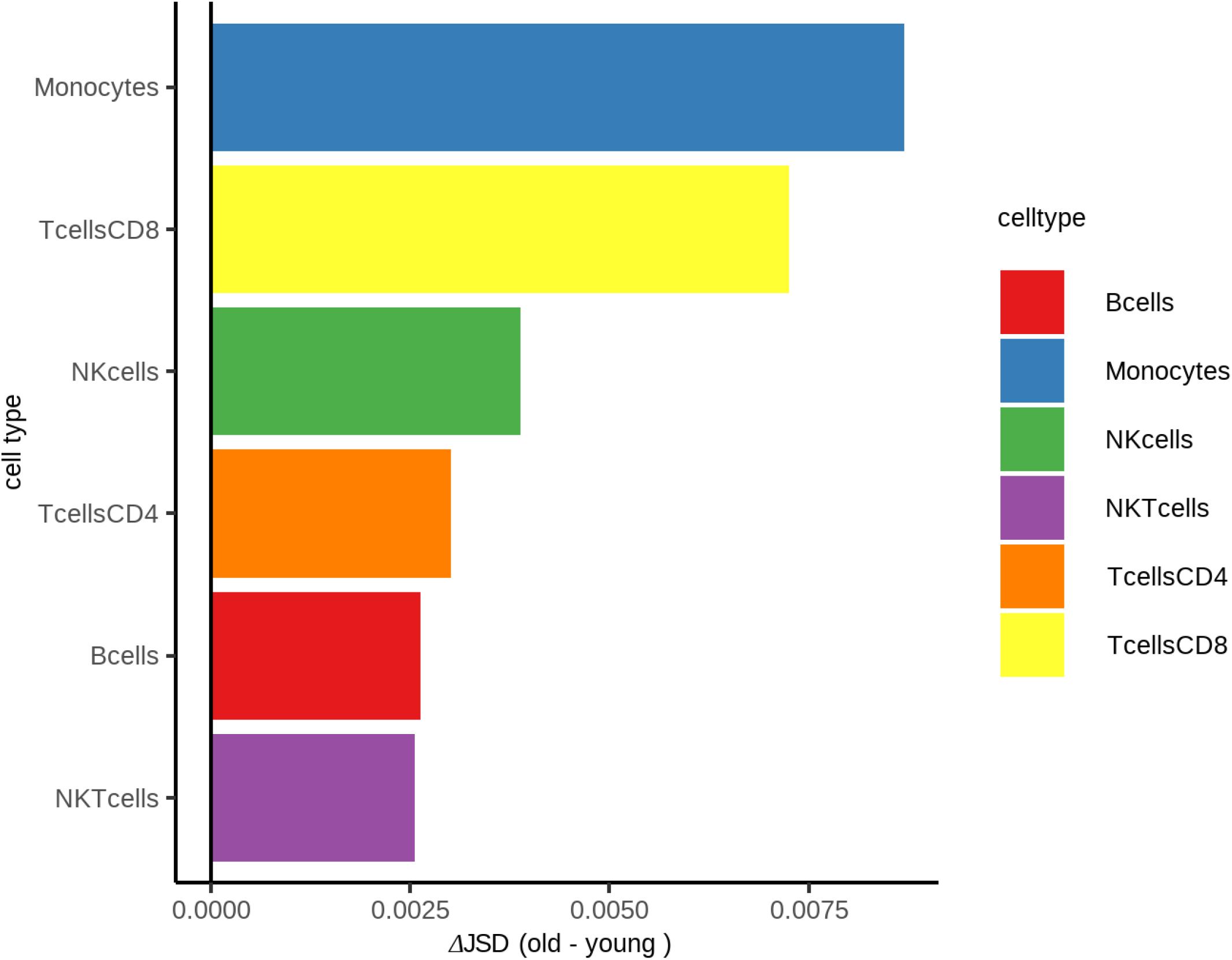
JSD applied to individual level cell expression estimates from GTEx Whole Blood with 2 bins. Individual level cell expressions are estimated from CIBERSORTx high resolution mode using all genes.

**Fig S 16.**
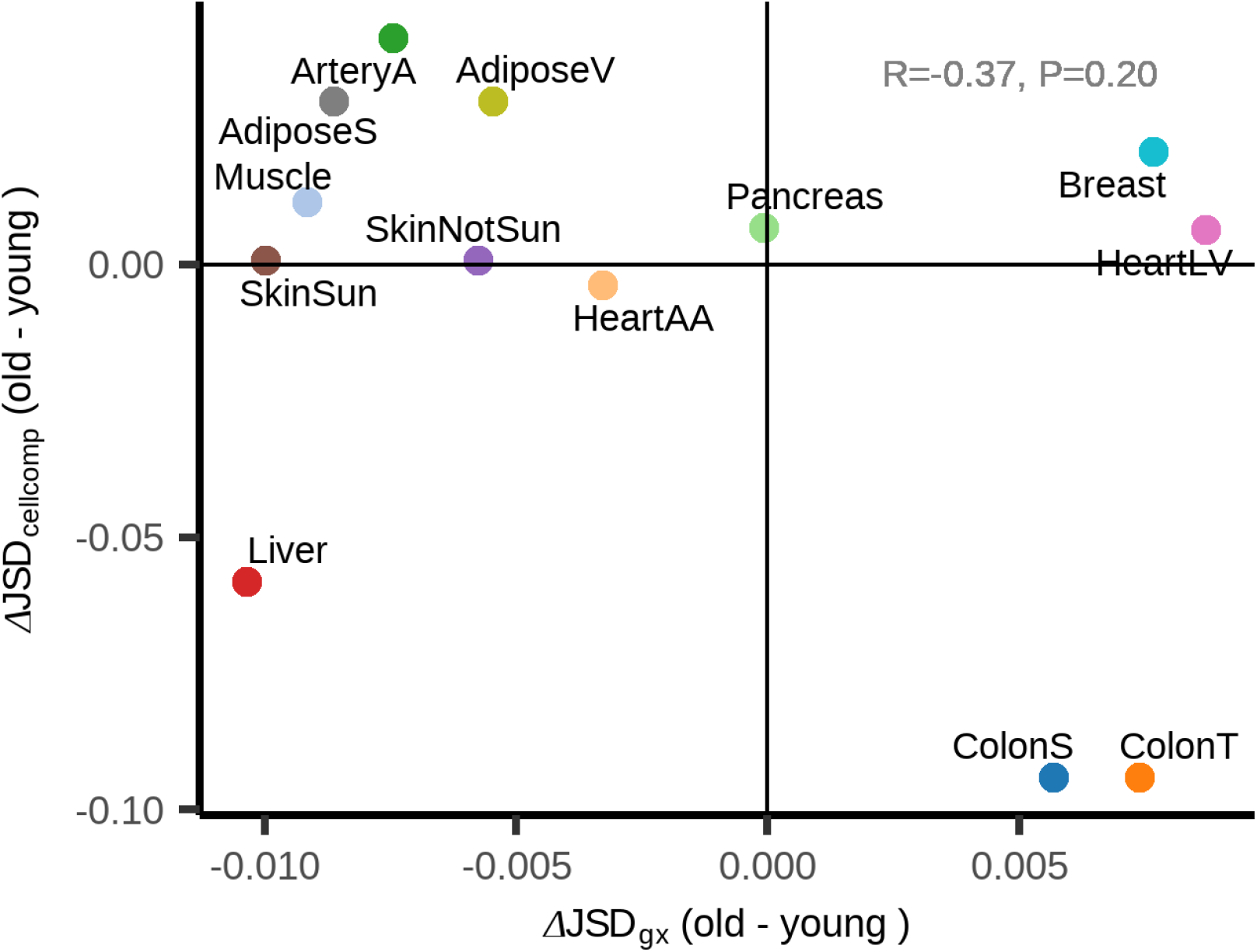
JSD heterogeneity metric on transcriptome profile with 2 age bins vs JSD applied to cell proportions. Cell proportion estimates are calculated using CIBERSORT. R and p-value are obtained from pearson correlation test.

**Fig S 17.**
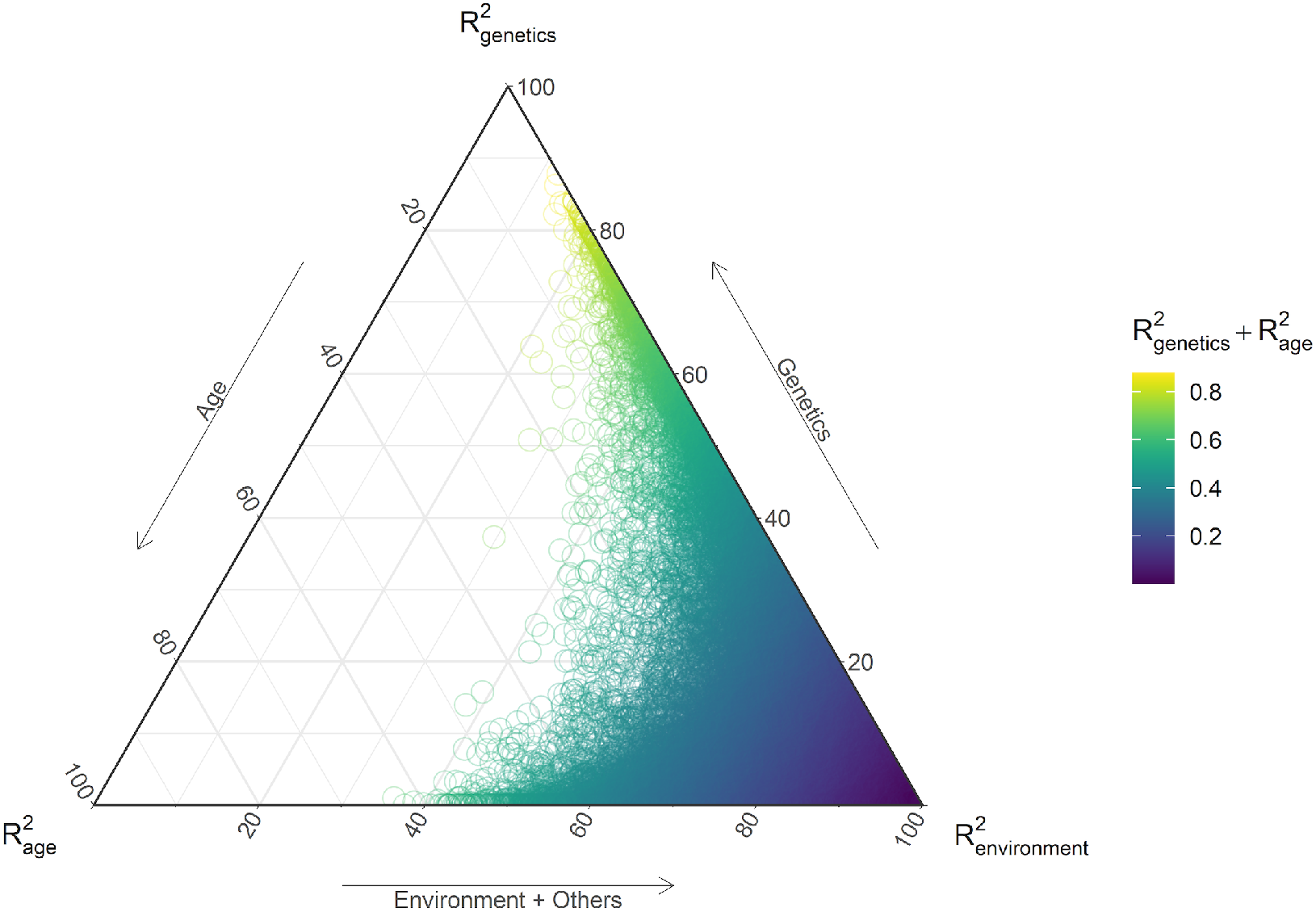
Ternary plot of the proportion of each genes expression variance explained by age, genetics and the environment.

**Fig S 18.**
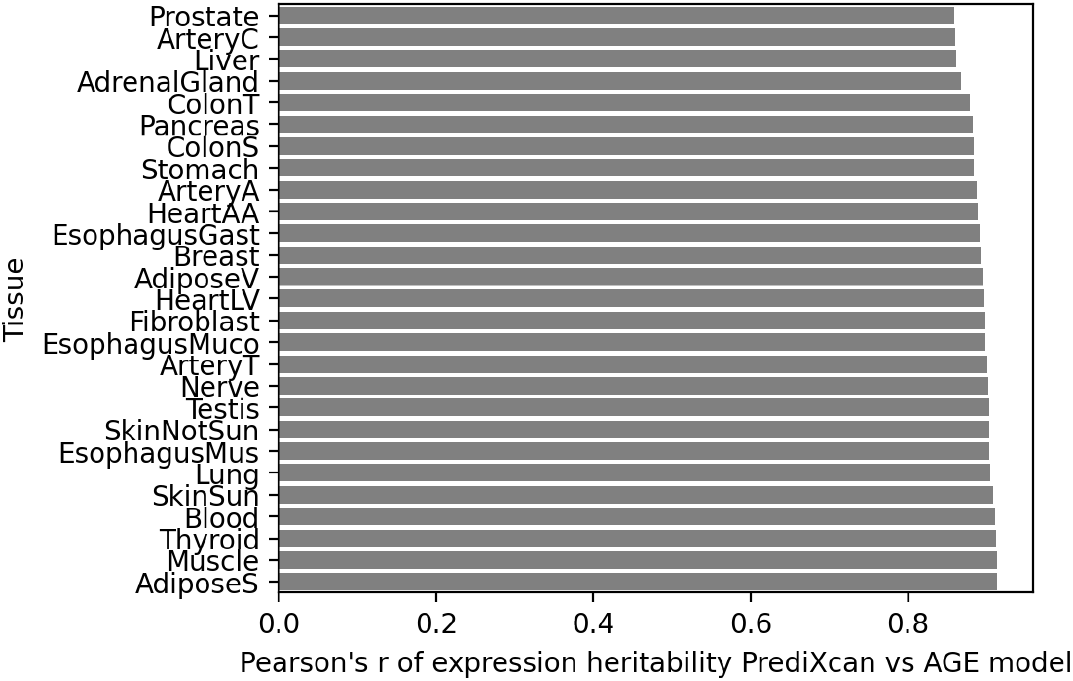
Pearson’s r of heritability estimate from PrediXcan (PredictDB) vs our model for each tissue.

**Fig S 19.**
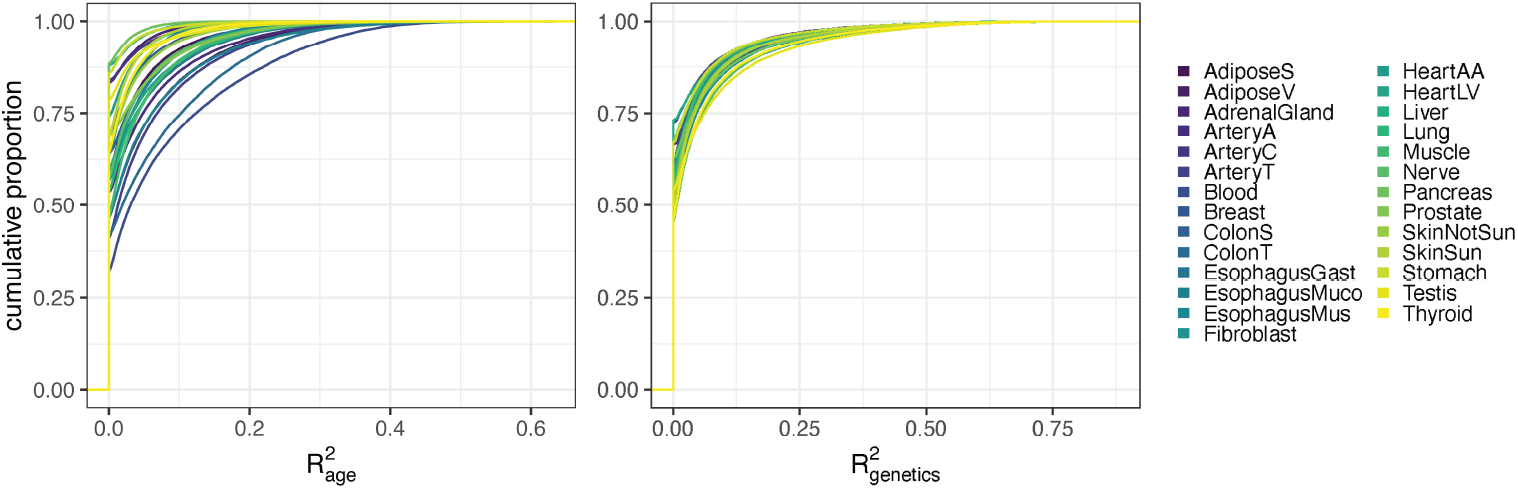
Cumulative distribution of 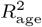 and *h*^2^ for all modeled genes within 27 tissues.

**Fig S 20.**
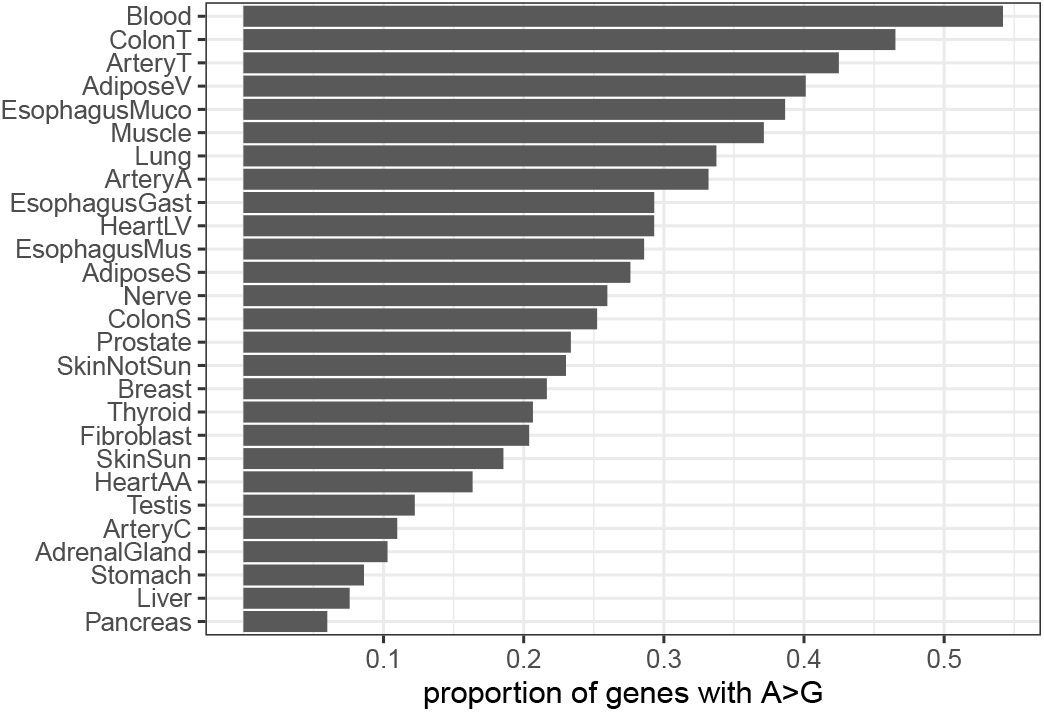
Proportion of genes within a tissue that have 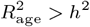.

**Fig S 21.**
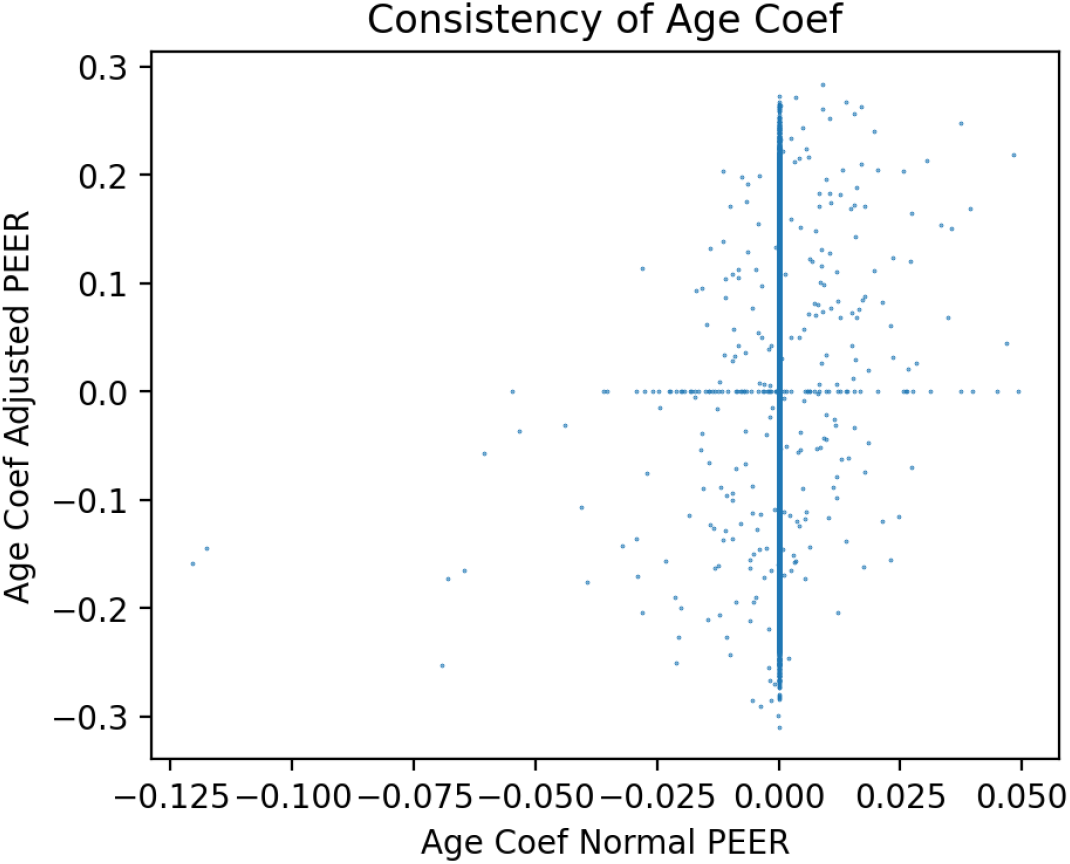
Scatter plot of each gene’s *β*_age_ of multiSNP model using GTEx PEER factors vs age-independent PEER factors for Whole Blood.

**Fig S 22.**
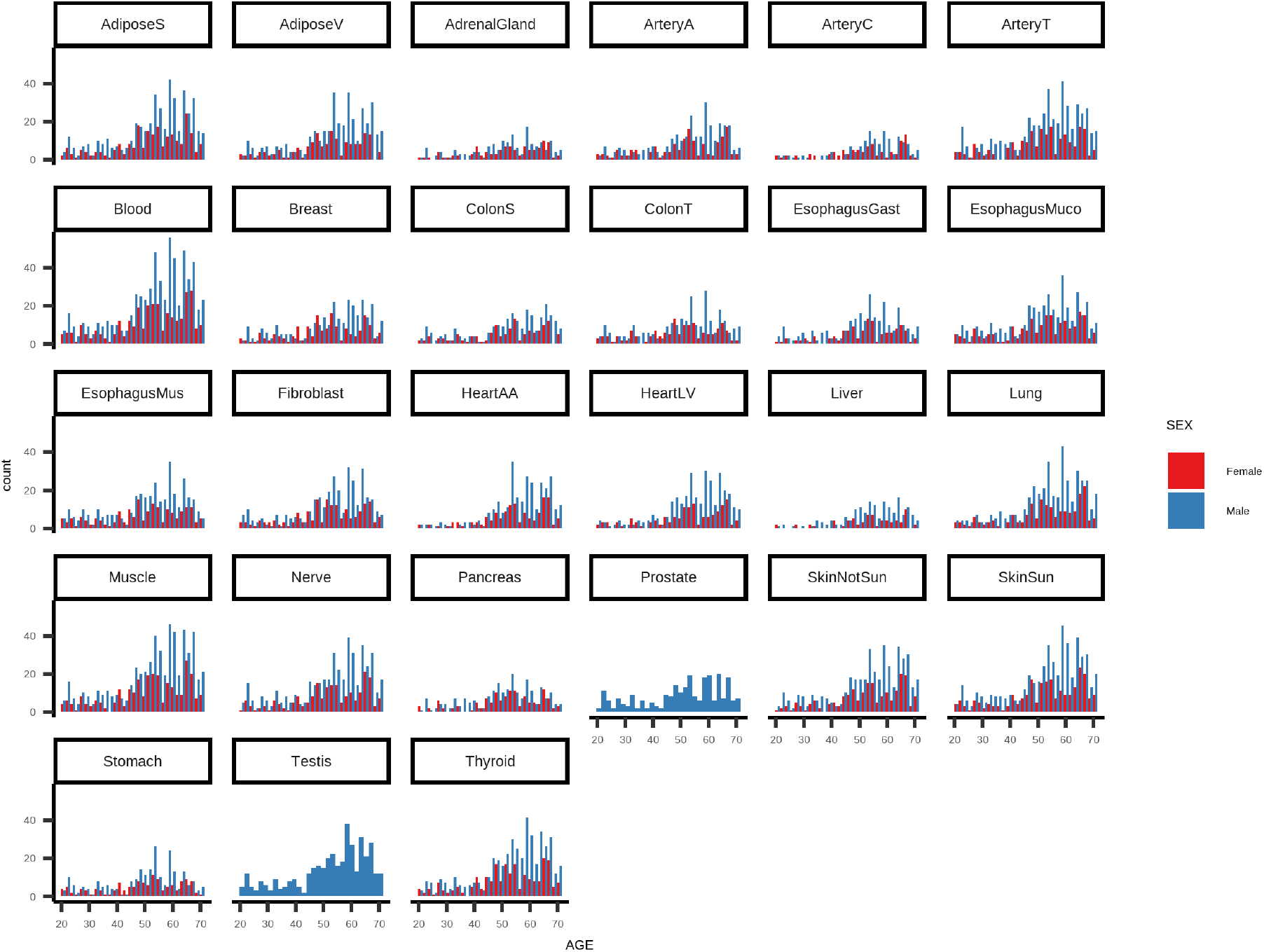
Age distribution in each tissue stratified by sex.

**Fig S 23.**
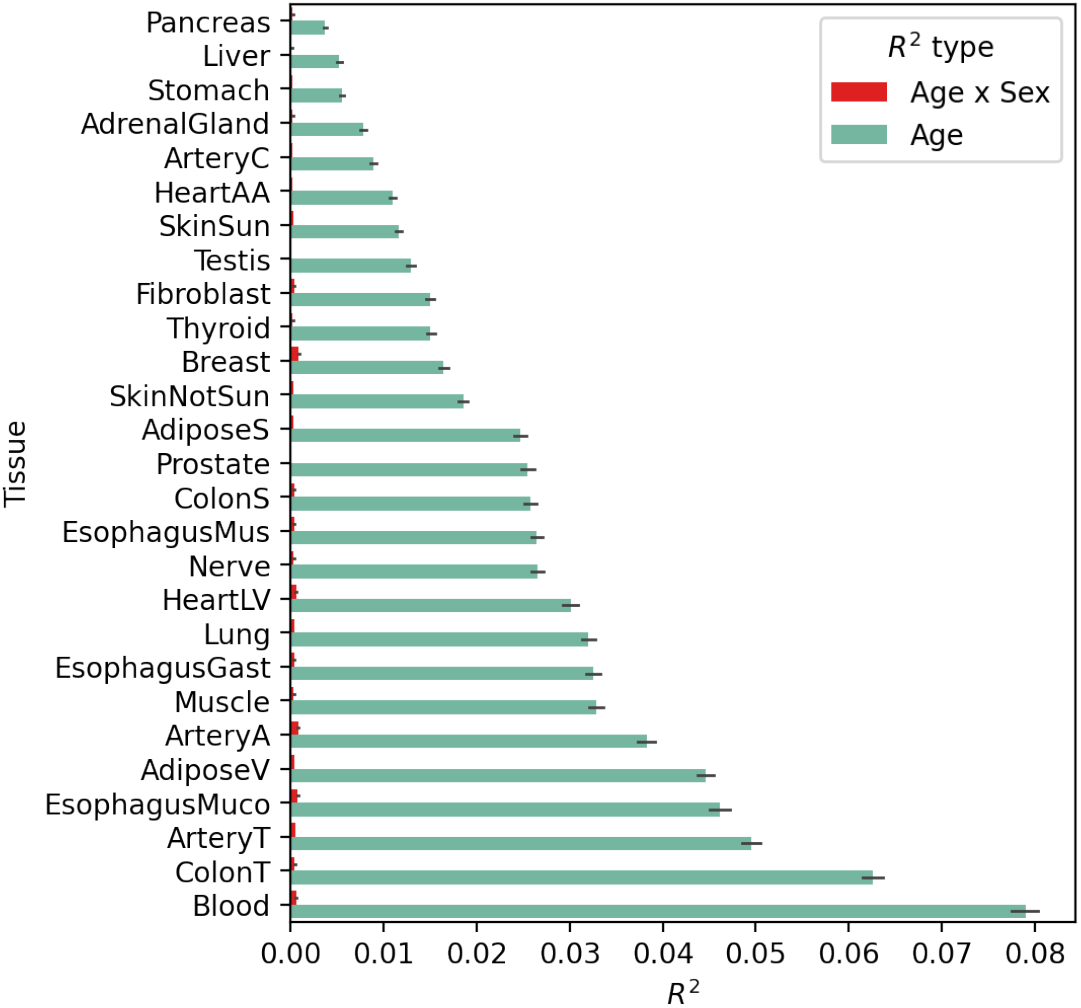
Average variance explained (*R*^2^) across all tissues for age term and age-sex interaction term.

**Fig S 24.**
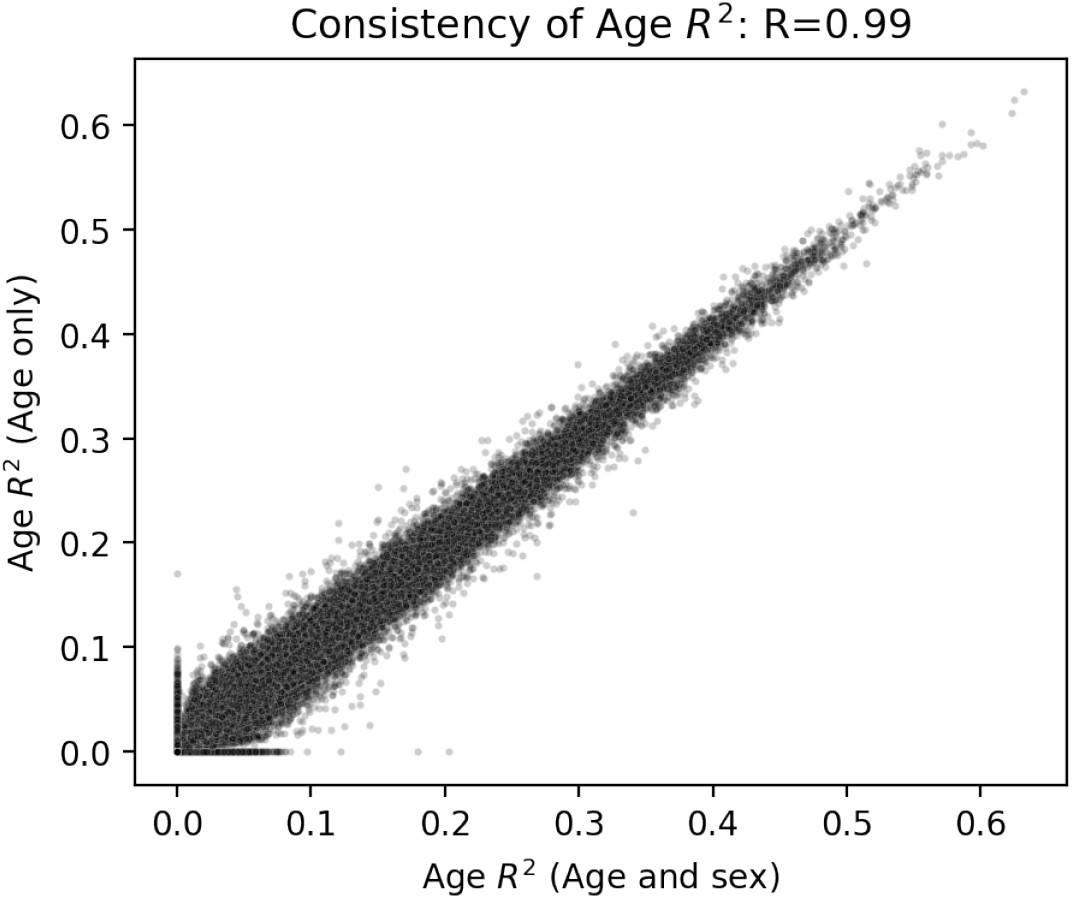
Variance explained (*R*^2^) across all genes and tissues for age term for joint age and genetics model and model that includes age-sex interaction term.

**Fig S 25.**
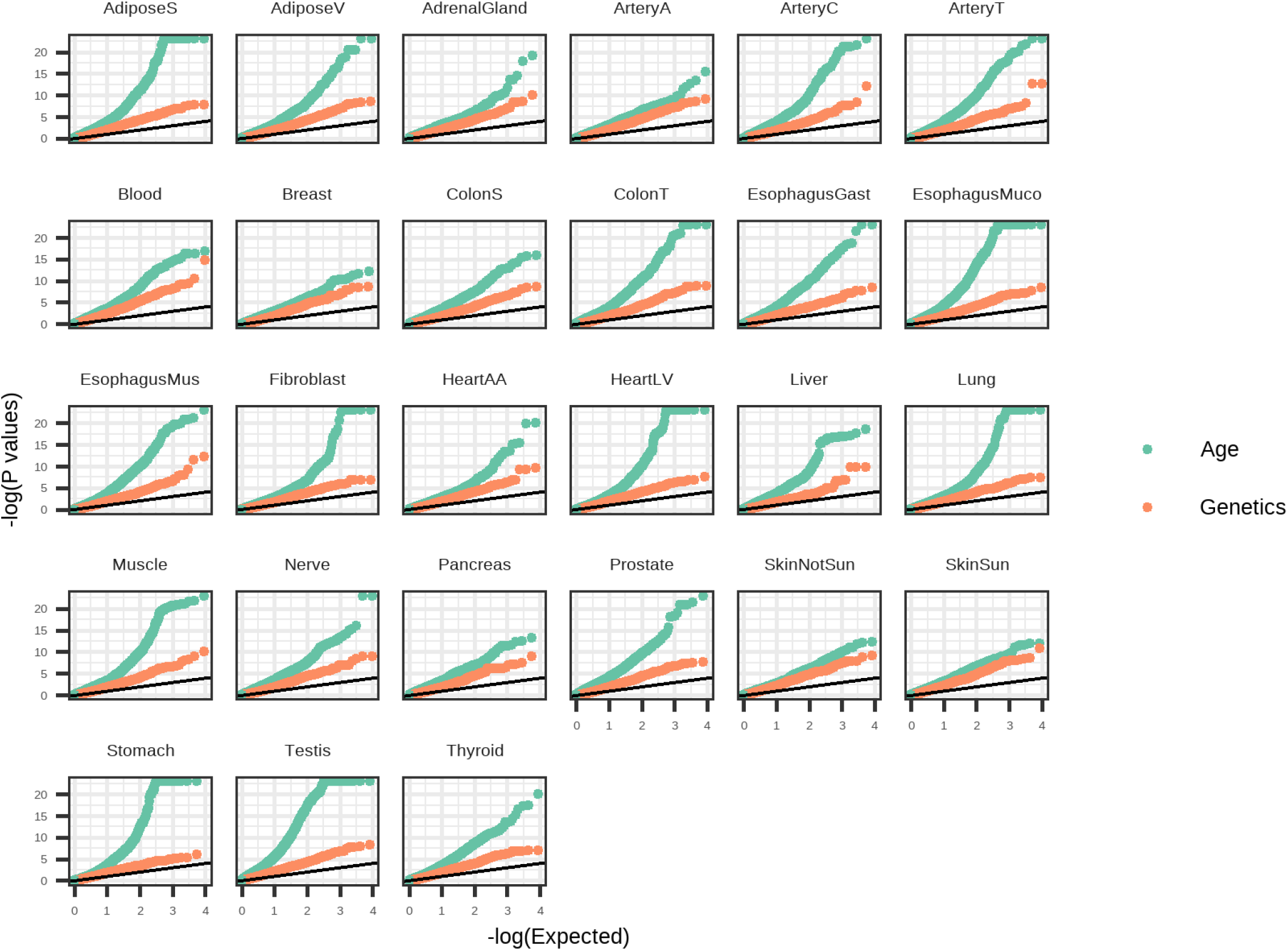
GO gene set enrichment P-values across all tissue for genes ranked by either *h*^2^ or 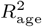.

**Fig S 26.**
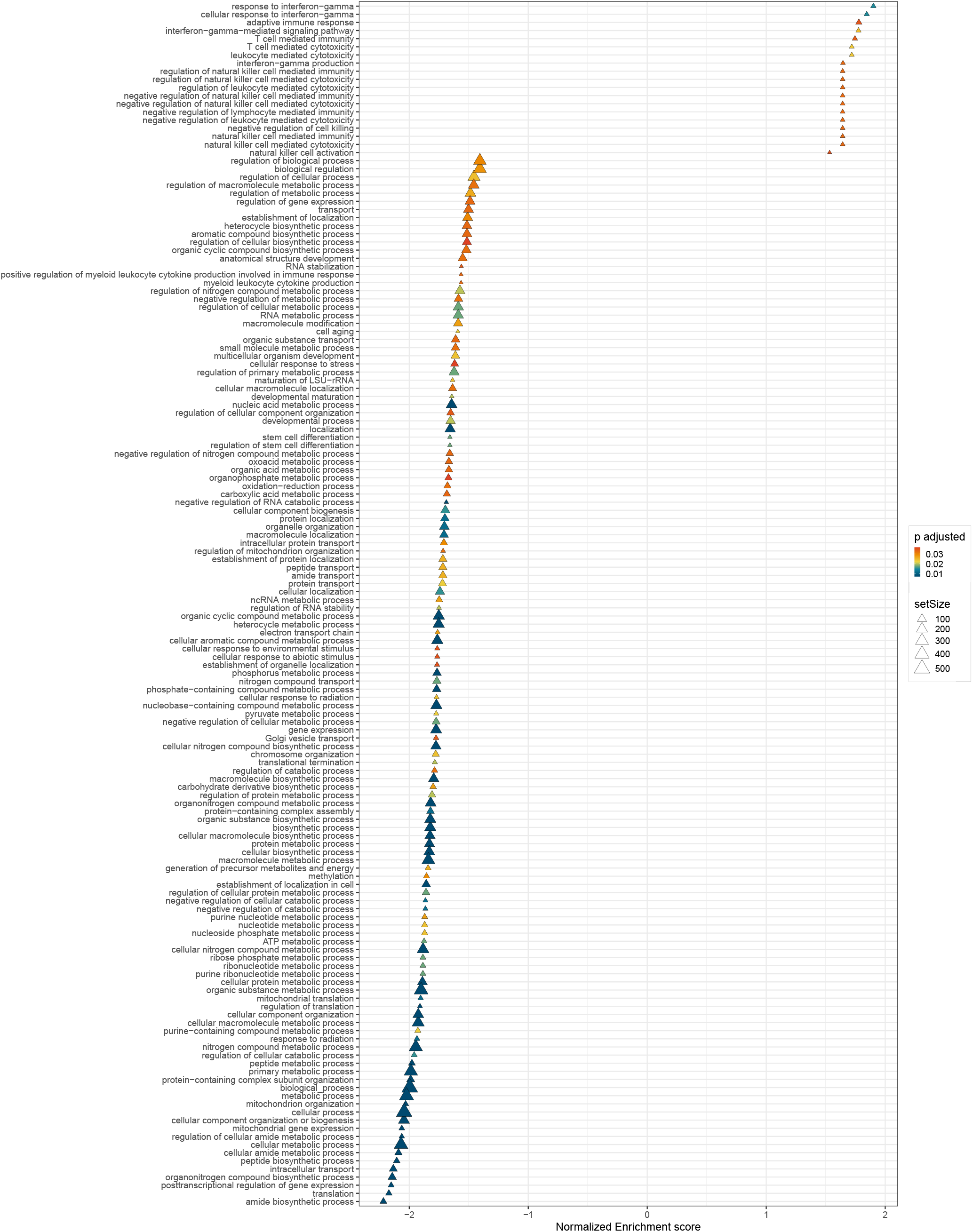
Pruned list of 50 GO Biological Processes identified in a GSEA using genes ranked by tissue-average *β*_age_ (FDR 0.05) (Full set of pathways in Table S5)

**Fig S 27.**
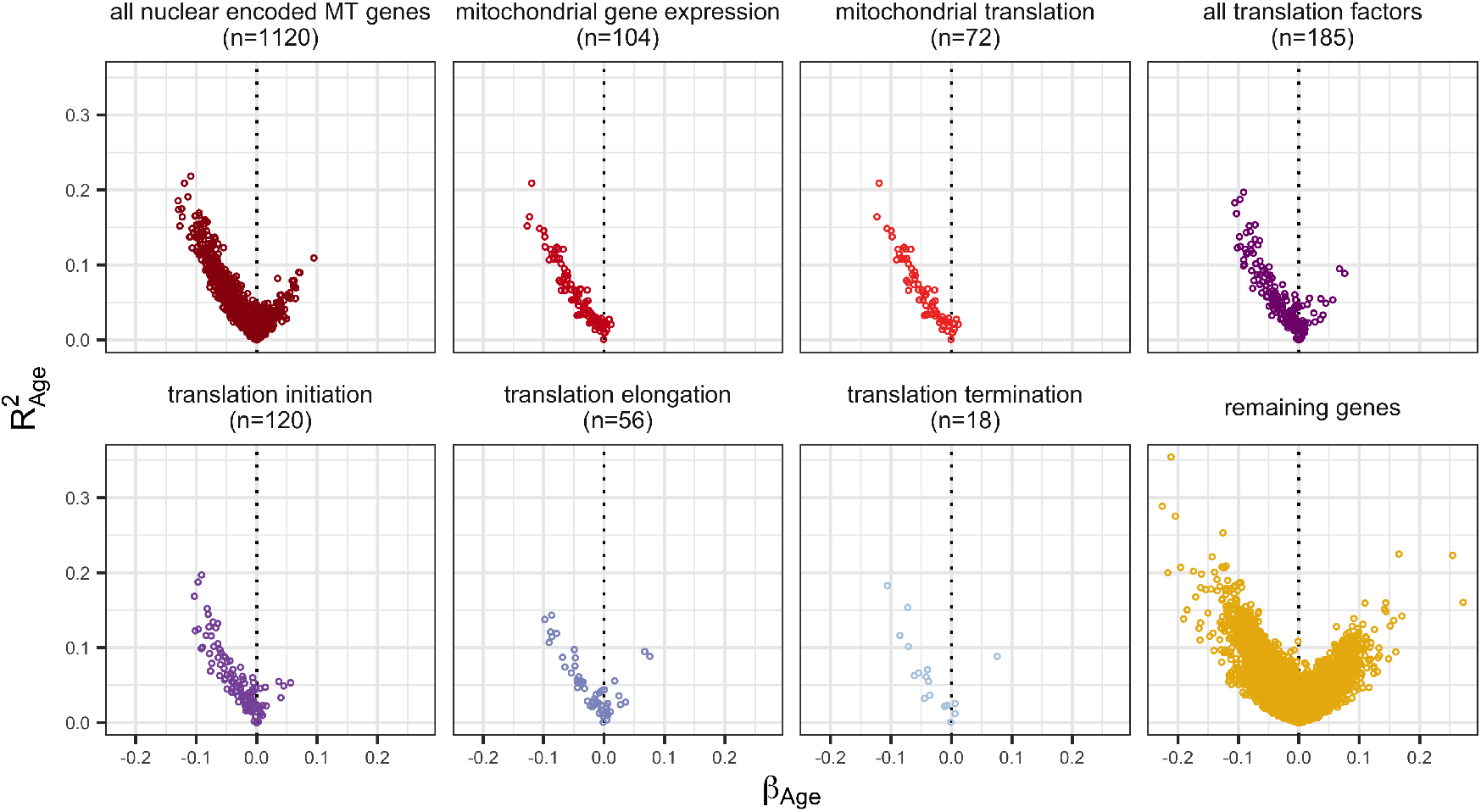
Distribution of tissue-averaged *β*_age_ and *R*^2^ for genes associated with specific mitochondrial and translation pathways.

**Fig S 28.**
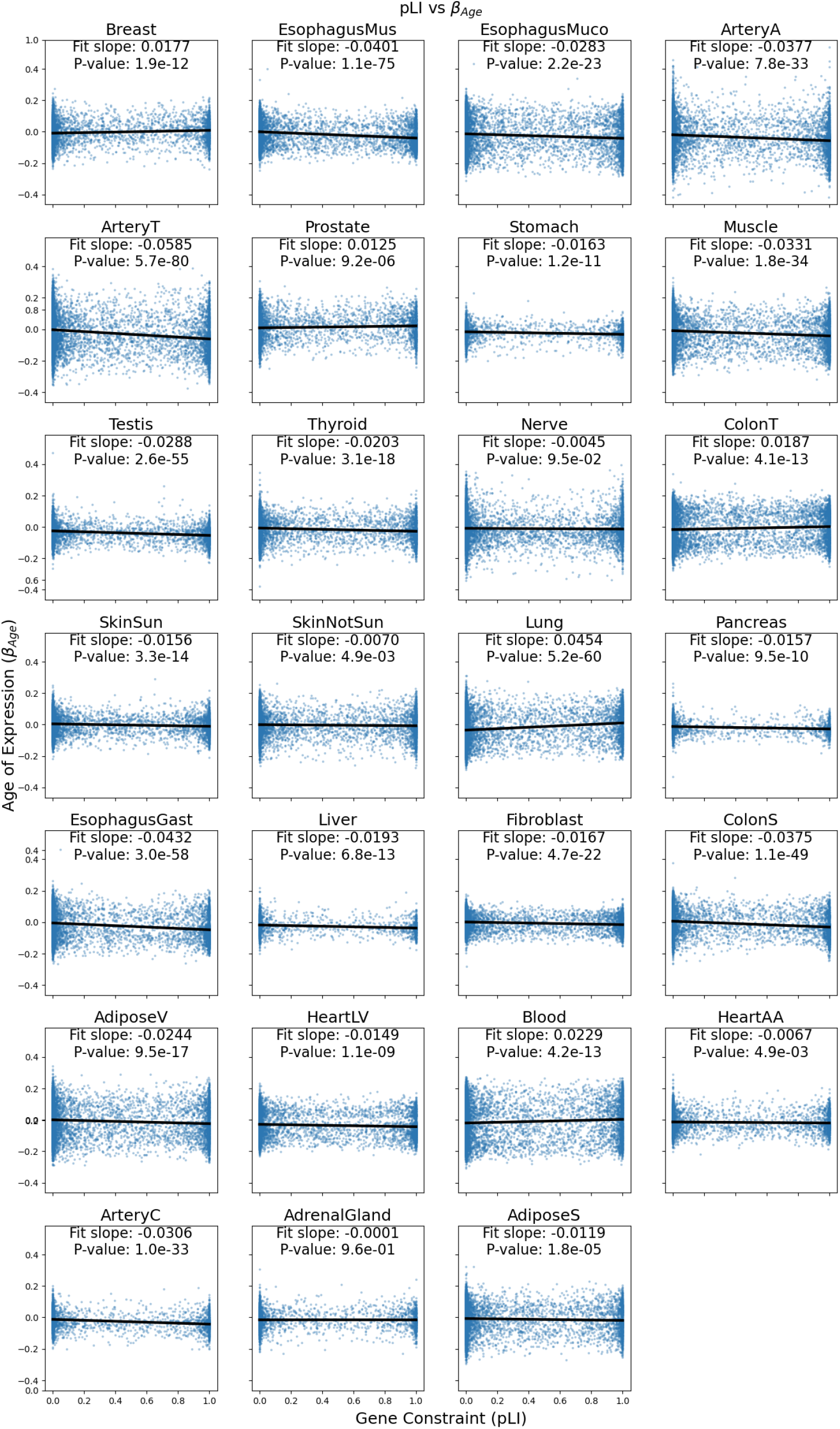
Age of expression (*β*_*Age*_) vs gene constraint (pLI) for all tissues with slope of linear fit and p-value of slope.

**Fig S 29.**
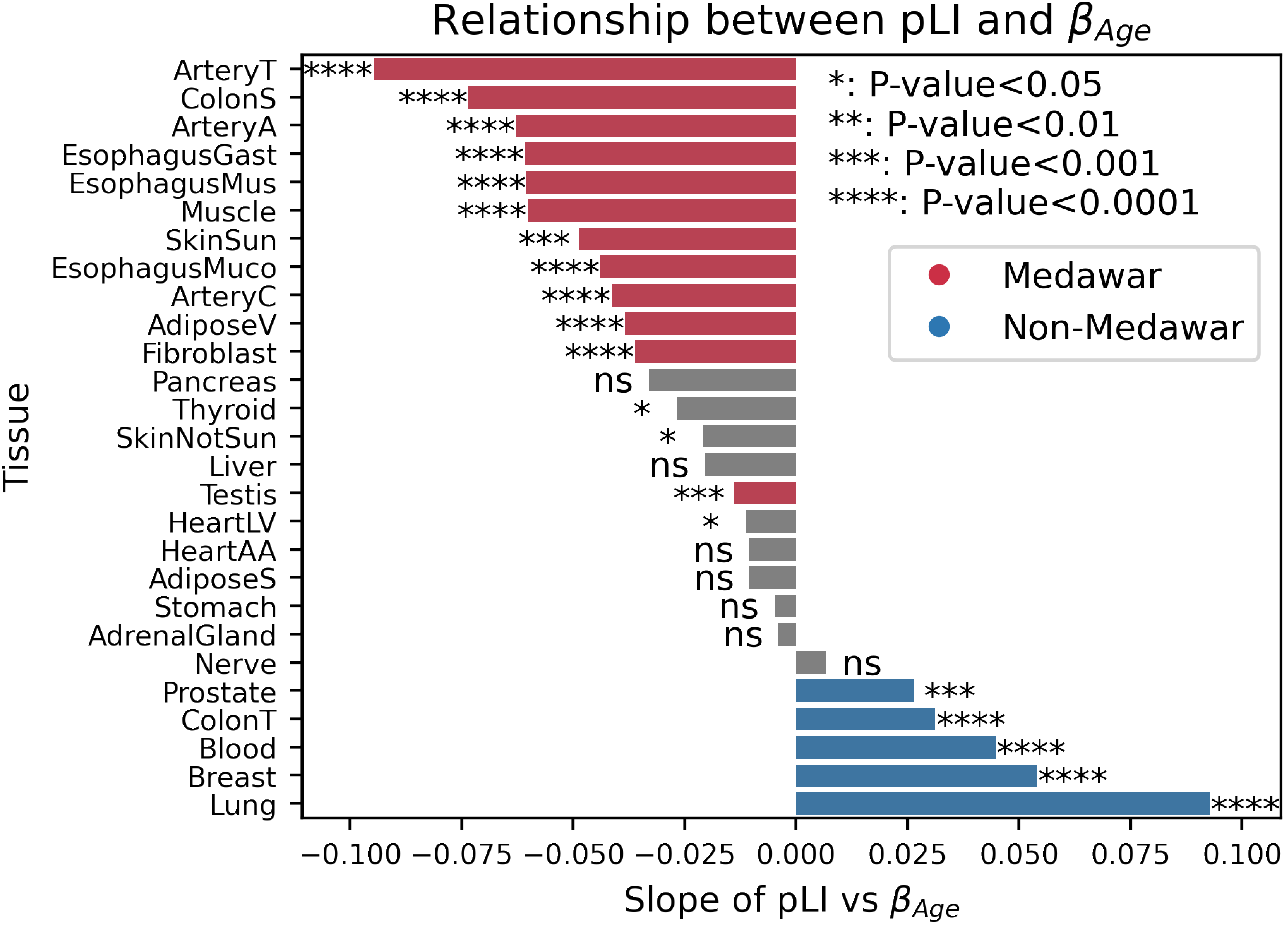
The slope of the relationship between gene constraint (pLI) and age of expression (*β*_*Age*_) across tissues for genes with 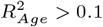. We repeated the analysis from figure 5D using only genes with ≥ 10% age-related gene expression variance. Color in this figure only indicates consistency with Medawar’s hypothesis with this subset of genes (Medawar - negative slope and P<0.001)

**Fig S 30.**
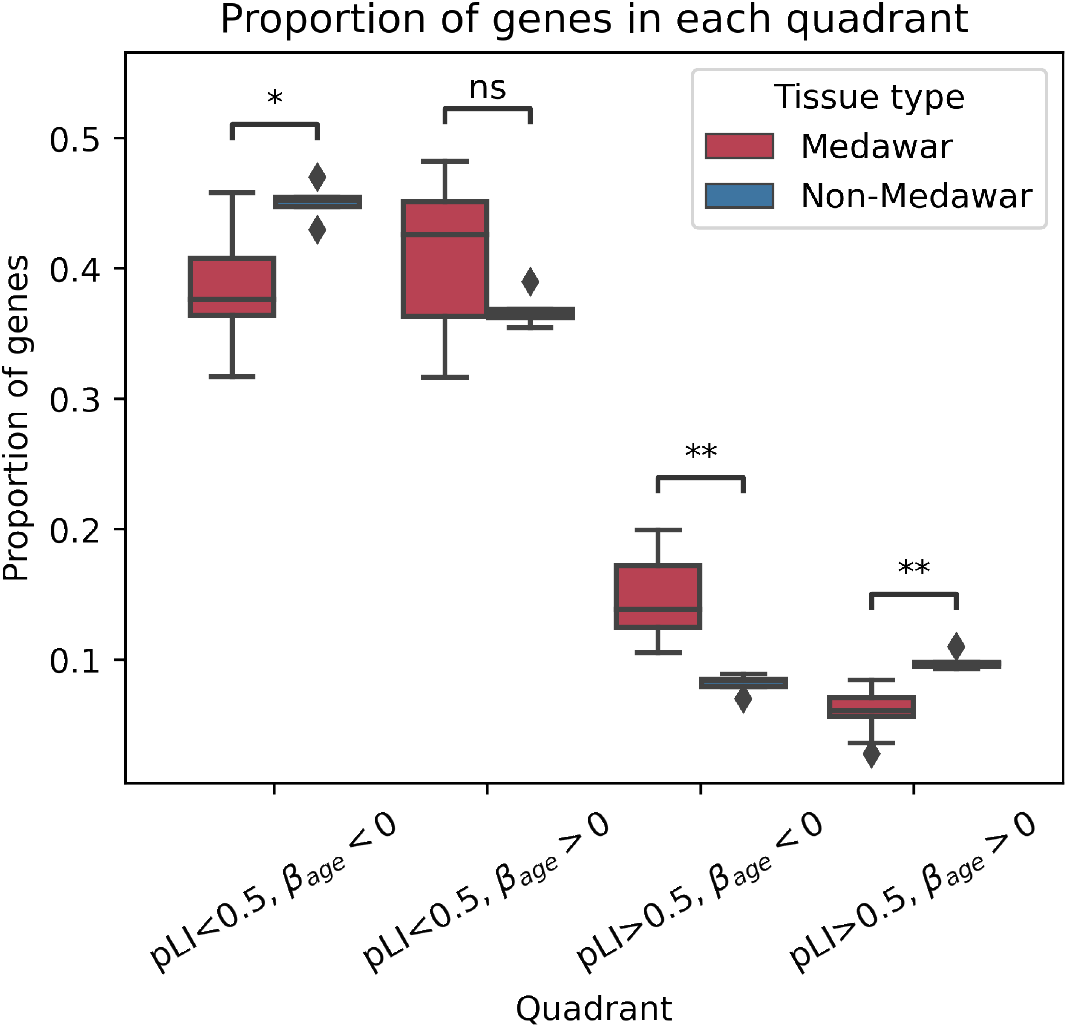
More highly constrained, late expressed genes in non-Medawar tissues than in Medawar tissues. We plot the proportion of genes within each quadrant of the gene constraint (pLI) vs. age of expression (*β*_age_) plots stratified by whether the tissue showed a significant Medawar or Non-Medawar trend. Center line of the boxplot indicates median, box limit indicates first and third quartiles, whiskers indicate the maximum and minimum and points indicate outliers.

**Fig S 31.**
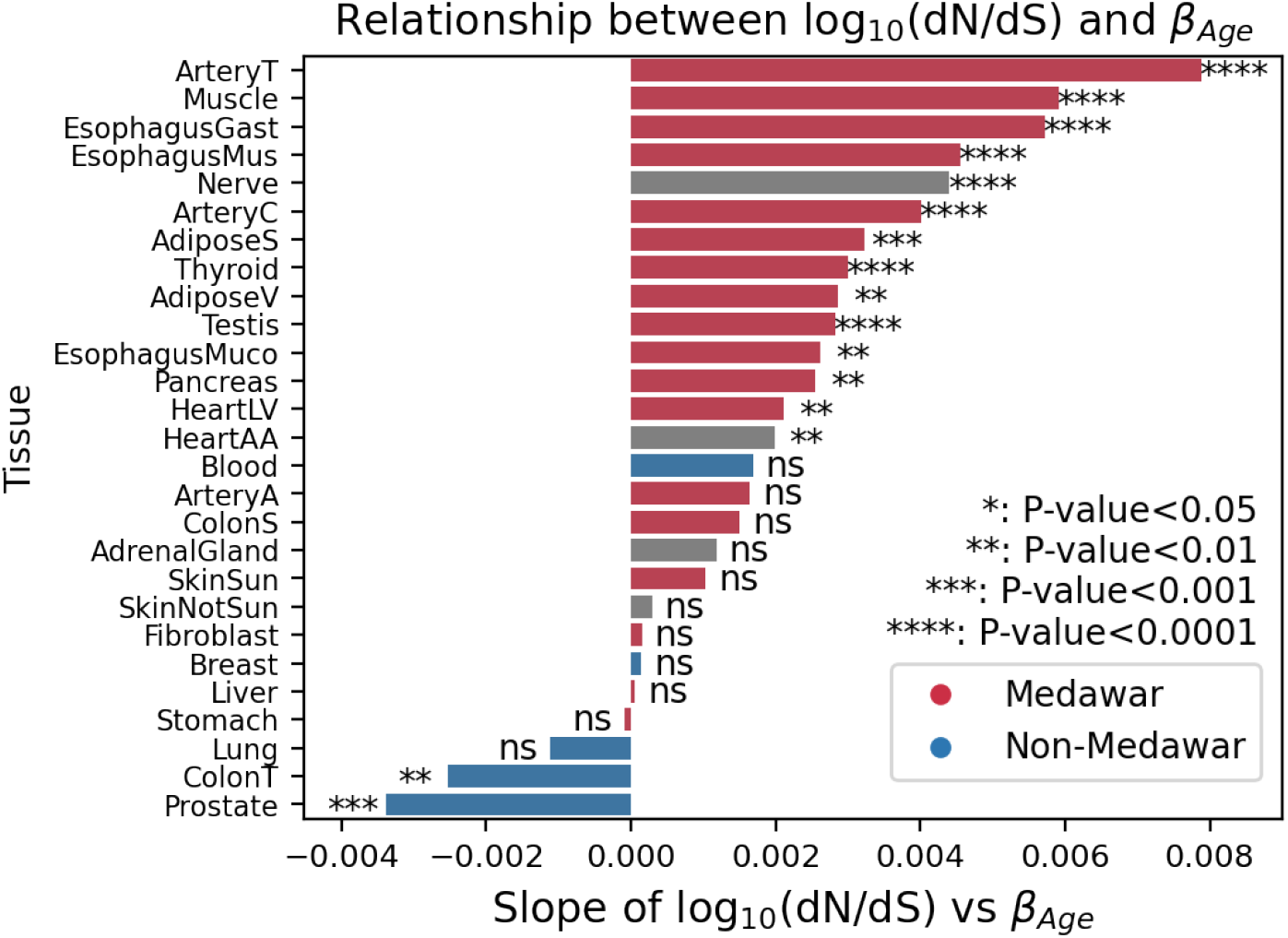
Linear regression slope of evolutionary constraint (dN/dS data from ortholog comparison between 8,175 human and chimpanzee genes) vs age-associated gene expression (*β*_age_) across tissues. Genes subsetted to 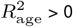.

**Fig S 32.**
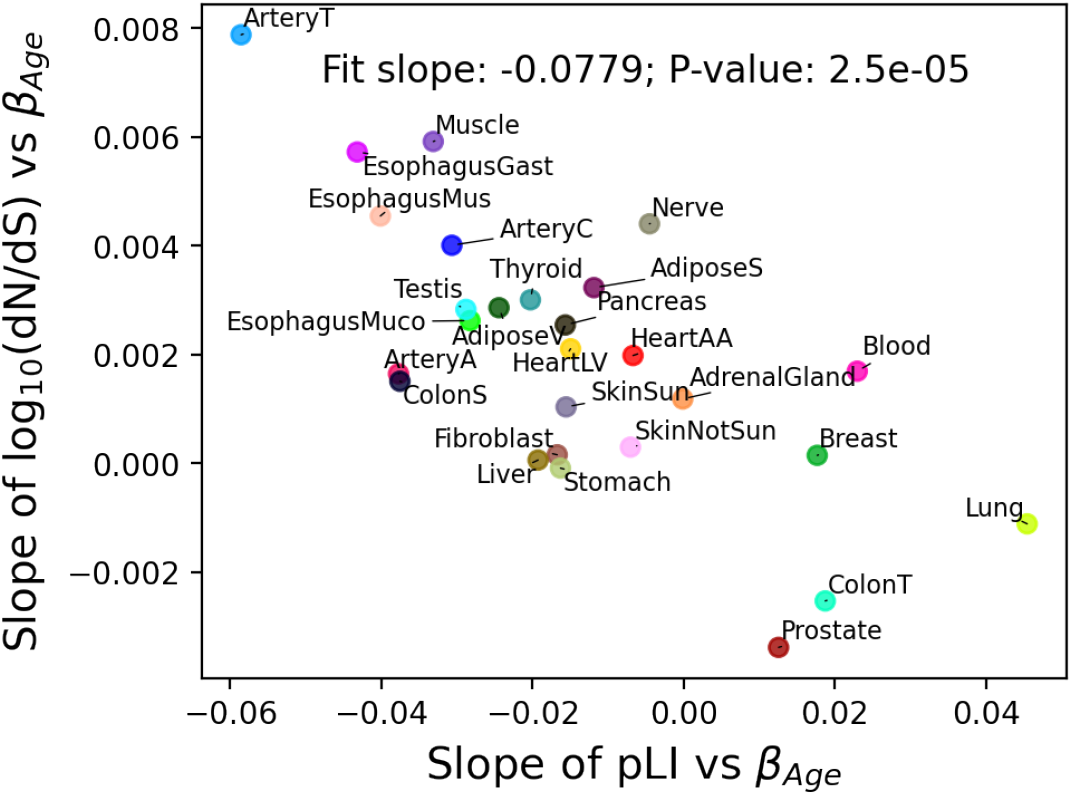
Consistency of Medawarian trend measures. We plot the slope of gene constraint metrics (pLI and dN/dS) vs age of expression (*β*_age_) for each tissue

**Fig S 33.**
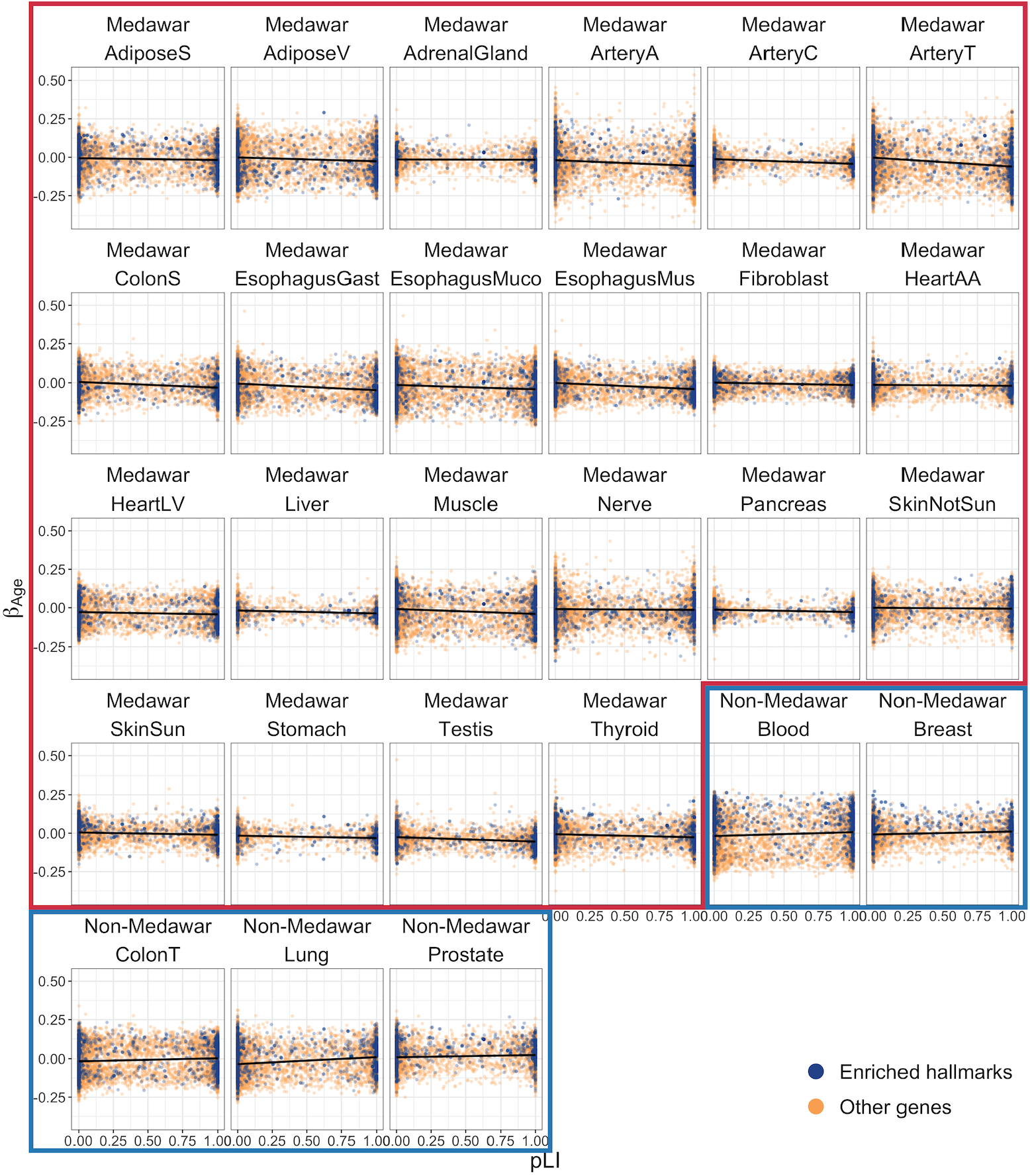
The relationship between *β*_age_ and pLI among tissues showing the strongest *Medawarian* and *non-Medawarian* signatures with genes in halmark pathways from (5D) highlighted in blue.

**Fig S 34.**
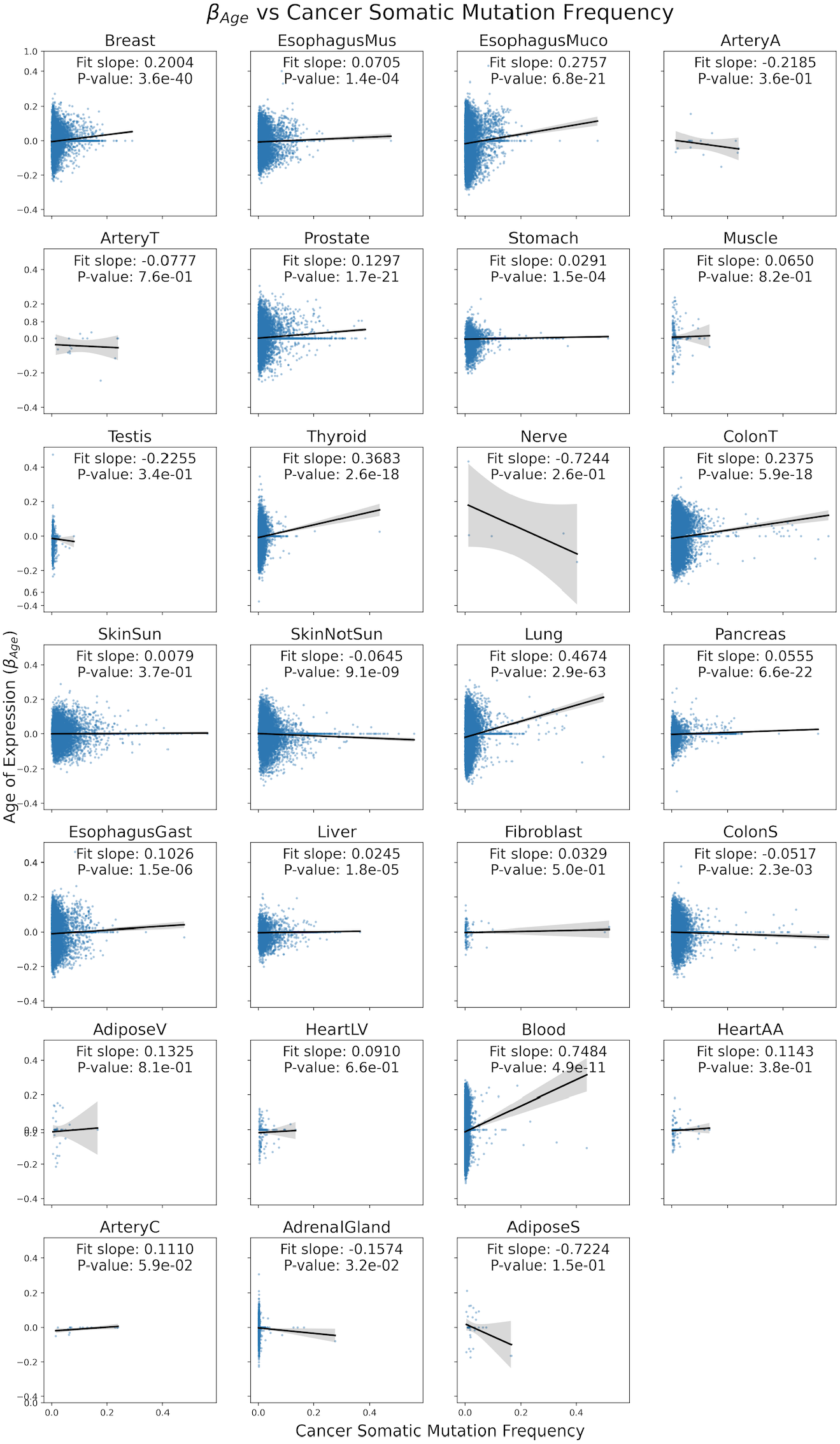
Age of expression (*β*_*Age*_) vs proportion of tumor samples with somatic mutation in gene across all tissues with slope of linear fit and p-value of slope. Genes included if they contain at least one sample with a somatic mutation within that gene.

**Fig S 35.**
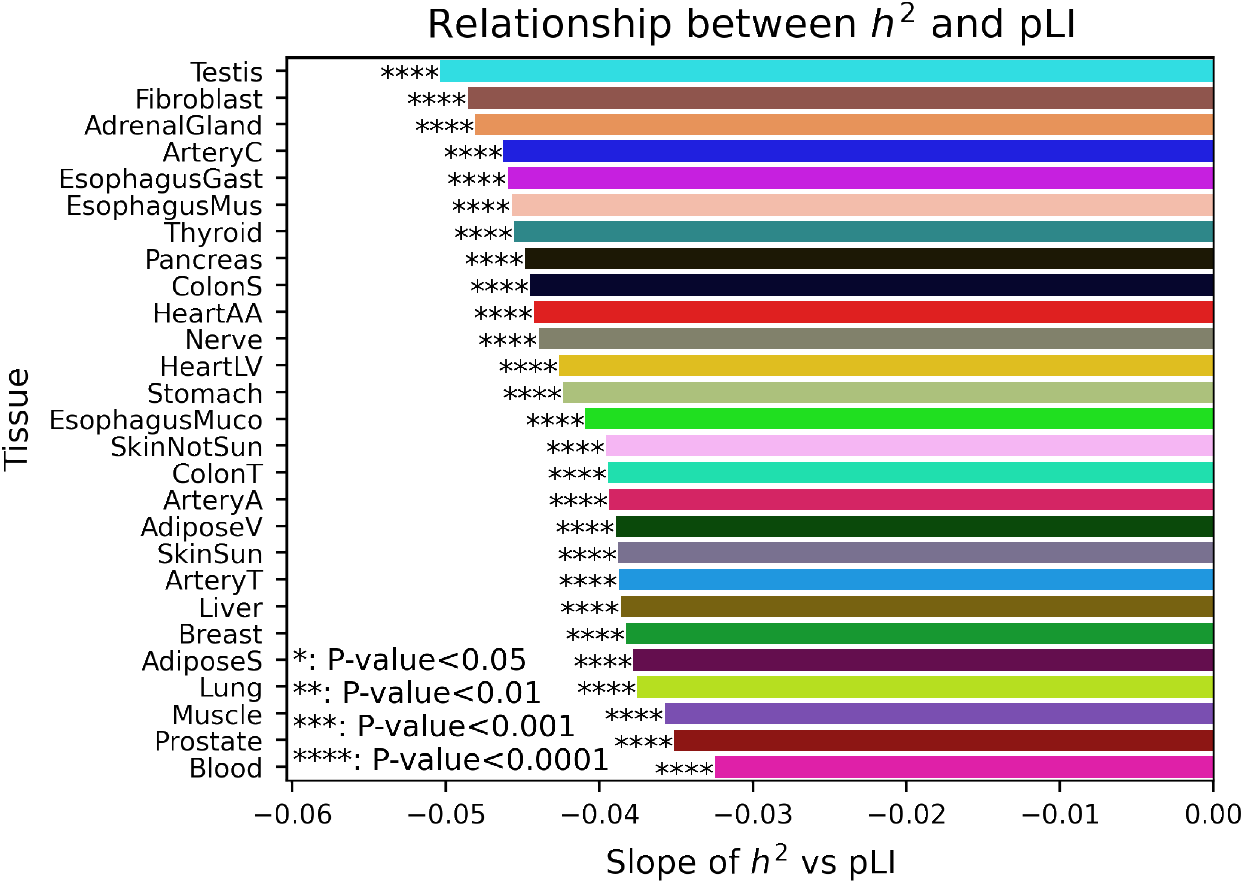
The slope of the relationship between constraint (pLI) and heritability (*h*^2^) across tissues. Slope and significance calculated from distribution of heritability and constraint for all tested genes (Fig. S36).

**Fig S 36.**
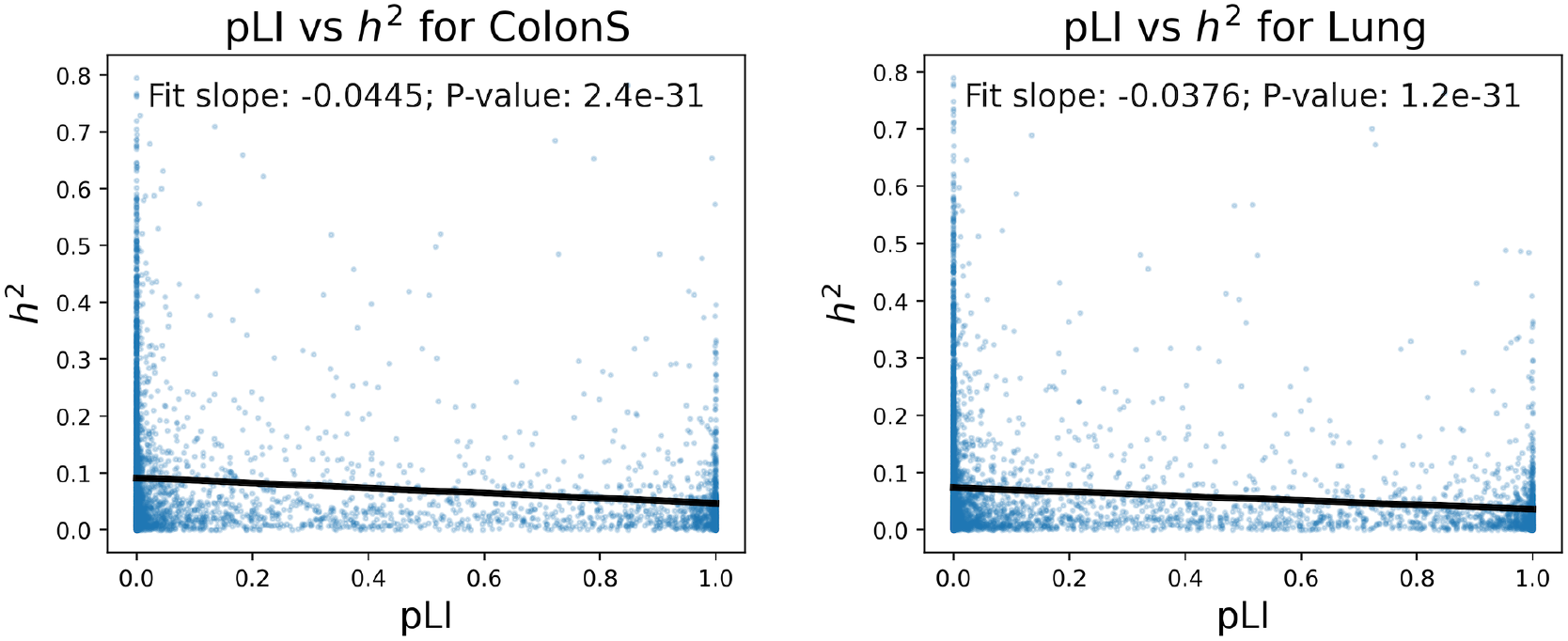
Distribution of gene expression heritability vs gene constraint (pLI) for all analyzed genes within two tissues. Slope and significance of relationship indicated.

**Fig S 37.**
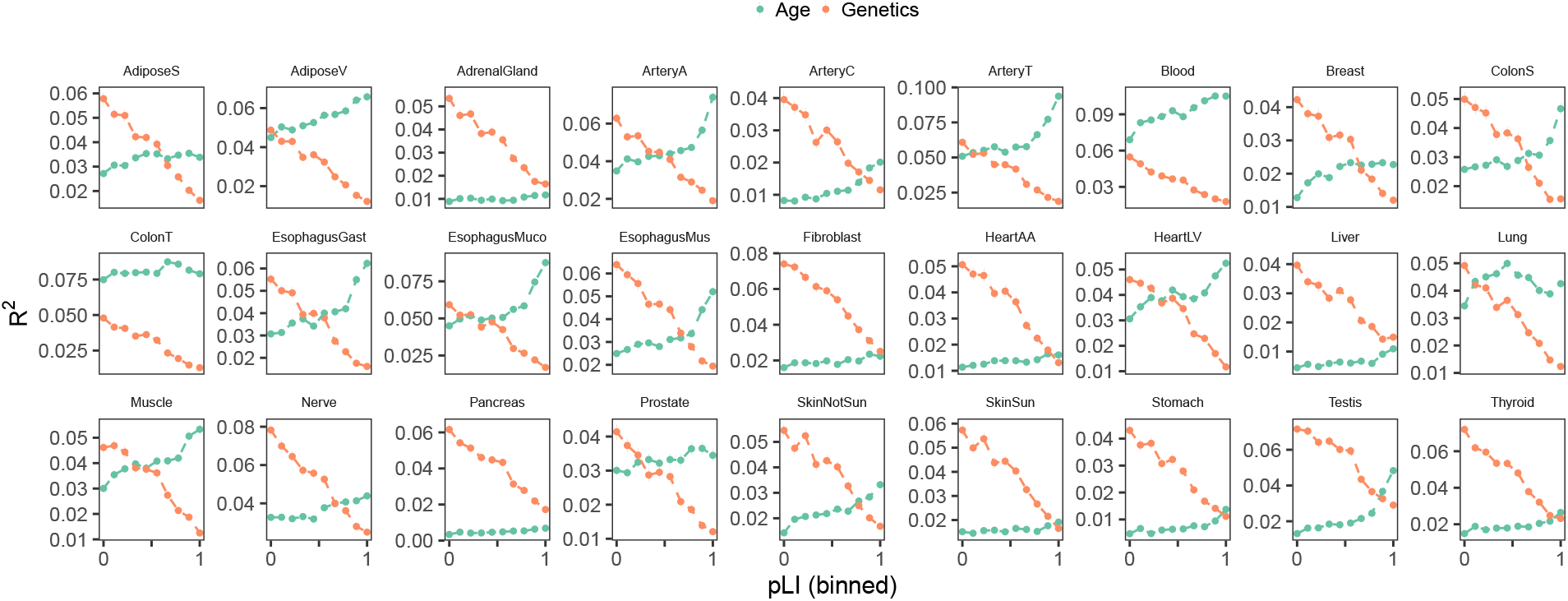
Relationship between gene expression variance explained by age and genetics and binned gene constraint (pLI) across tissues.

**Fig S 38.**
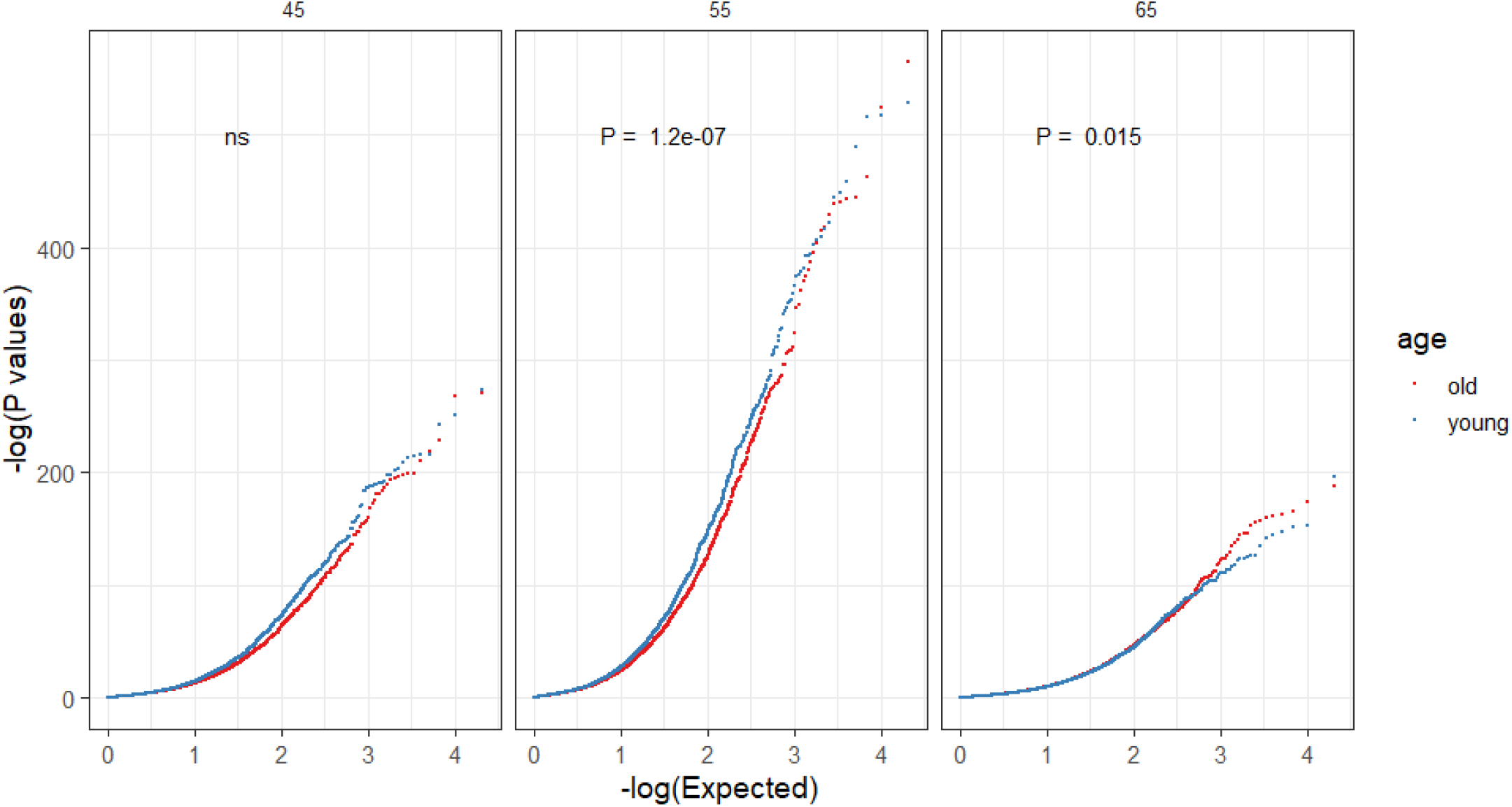
QQ plot of eQTL p-values from GTEx Whole Blood using different age cutoffs for two age bins, 45, 55 and 65 years old respectively. P-values for distribution difference are obtained from two-sided Welch’s t test.

**Fig S 39.**
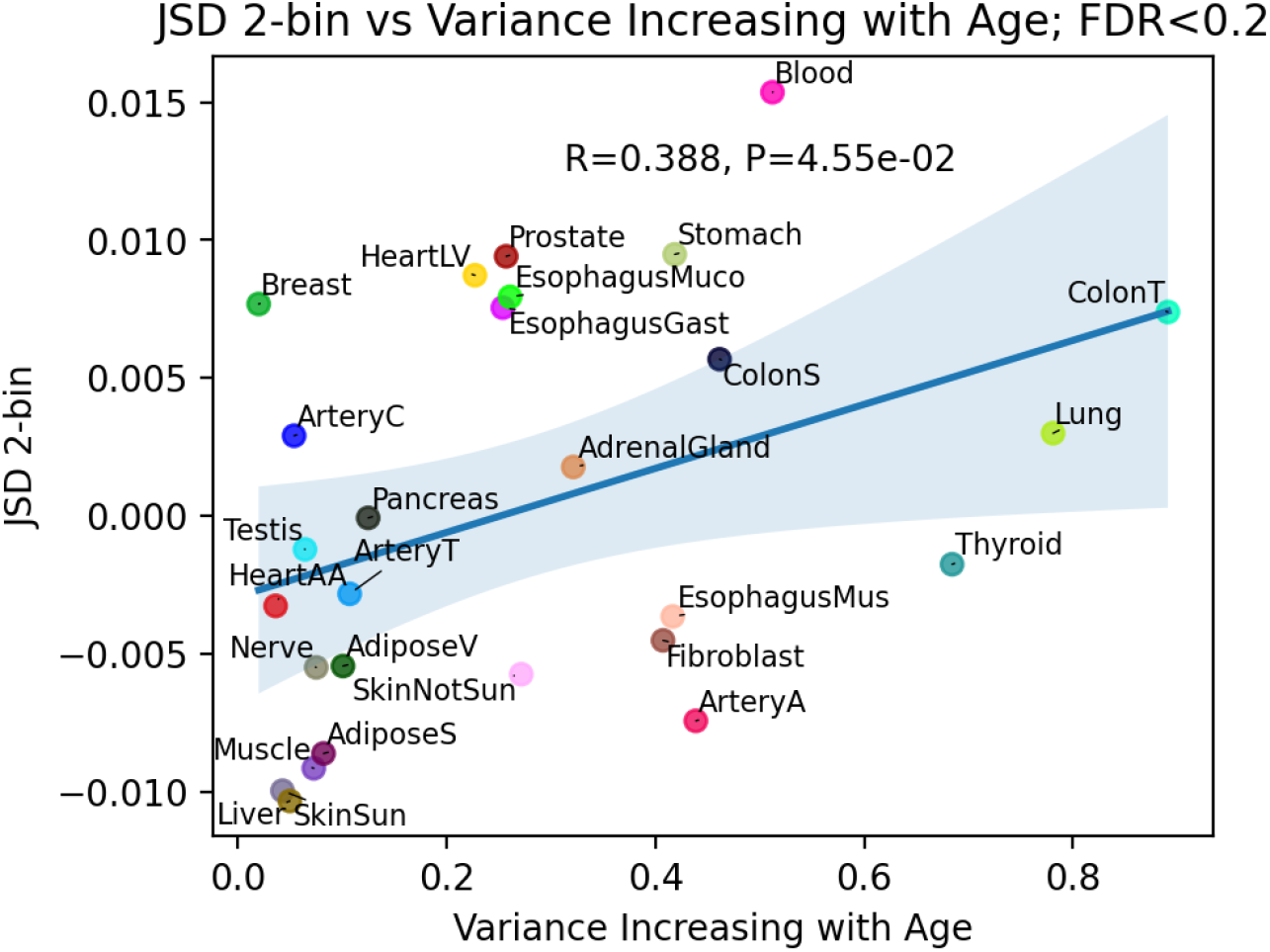
JSD age-related heterogeneity metric with 2 age bins vs proportion of heteroskedastic genes with increasing expression variance with age.

**Fig S 40.**
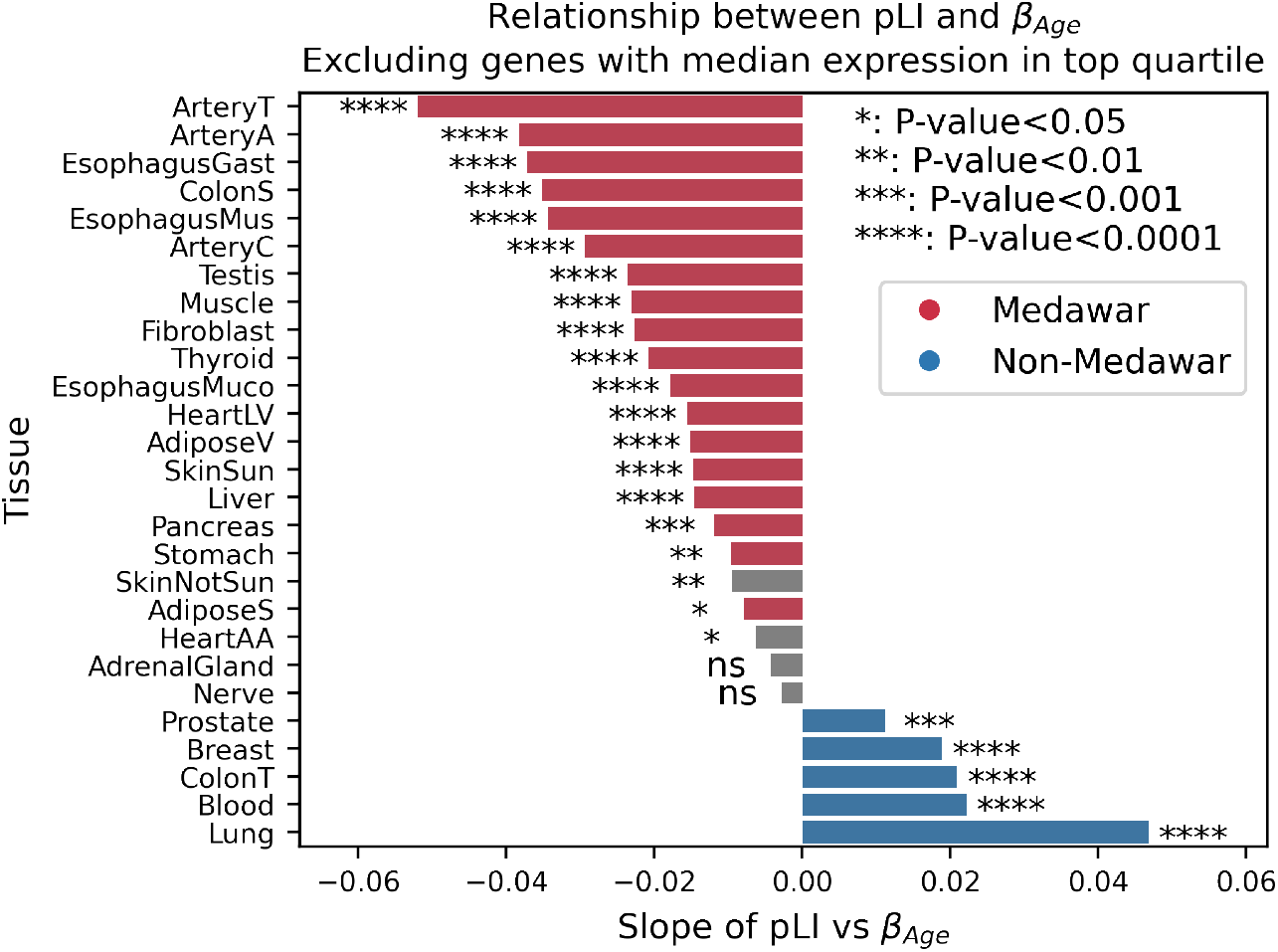
The slope of the relationship between gene constraint (pLI) and age of expression (*β*_*age*_) across tissues excluding highly expressed genes. We repeated the analysis from figure 5D using genes from the lower 3 quartiles of median gene expression from GTEx. Color indicates consistency with Medawar’s hypothesis from figure 5D. Spearman’s *ρ* 0.98 between slopes in 5D and this figure.

**Fig S 41.**
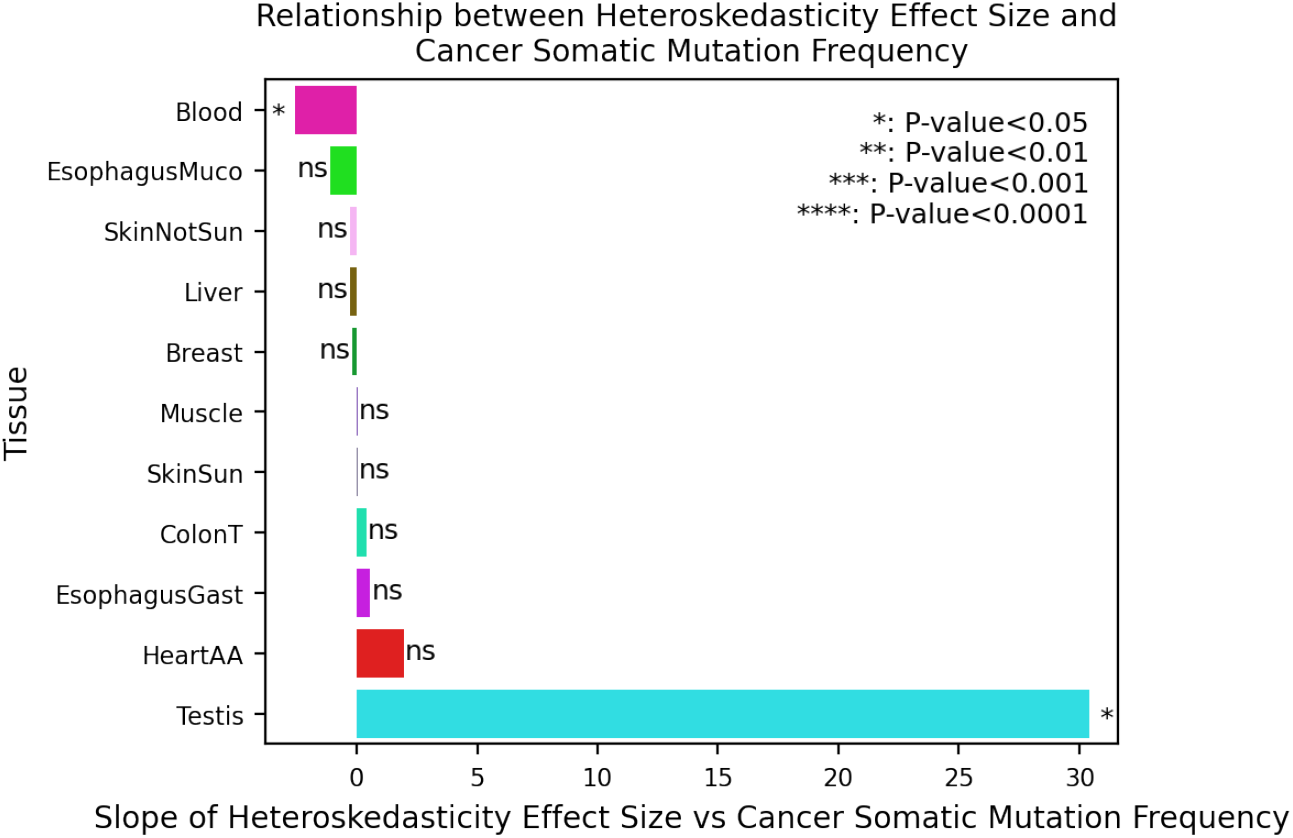
Effect size of age-related heteroskedasticity vs cancer somatic mutation frequency. Positive relationship indicates that genes with higher expression variance in old have higher rate of cancer somatic mutations. Only genes with >200 sequenced tumor samples in COSMIC and significant heteroskedasticity (FDR<0.2) are used. Tissues with *≤*5 genes meeting criterion are not show.

**Fig S 42.**
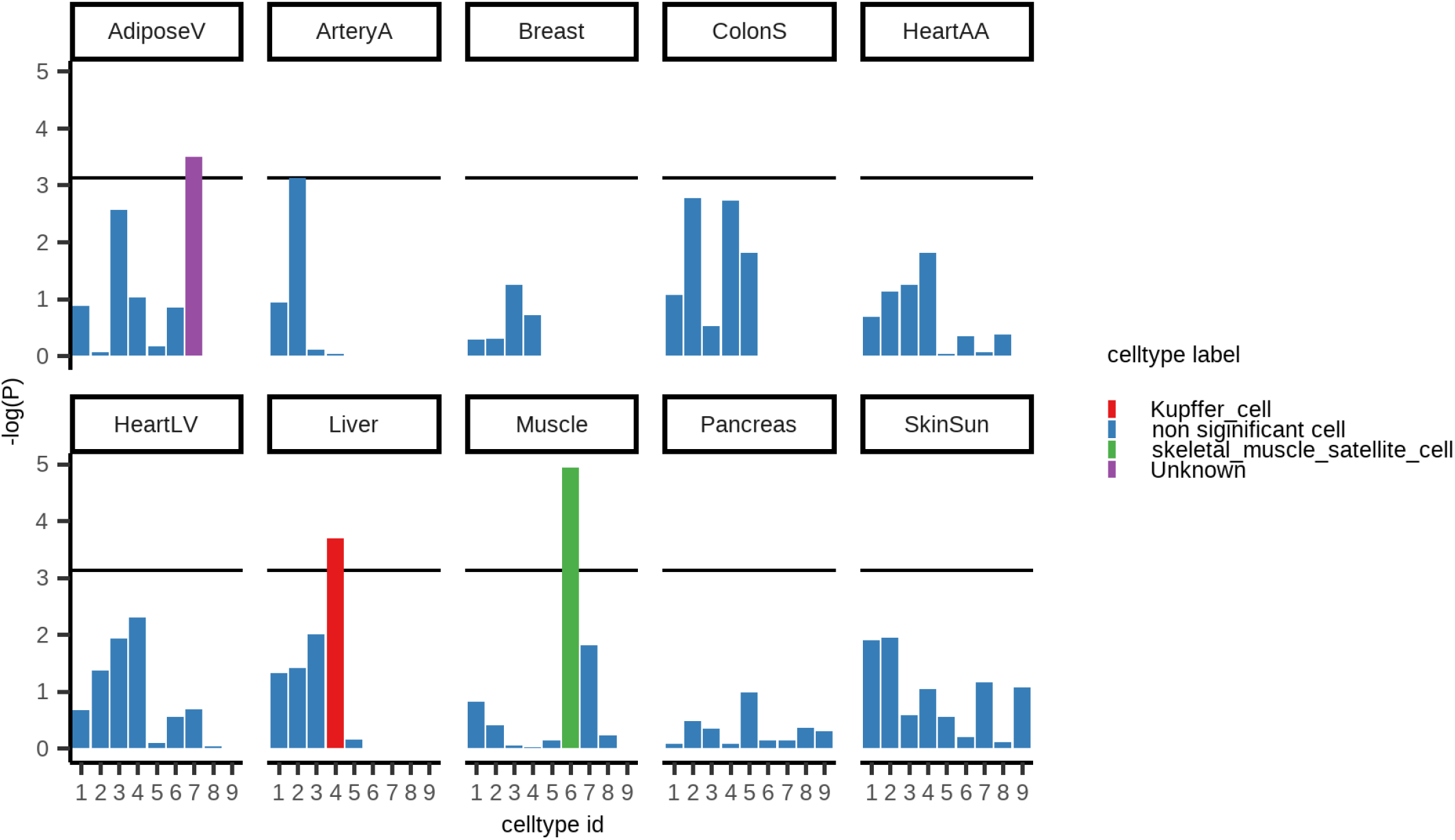
Breusch-Pagan heteroskedasticity test P-value for each cell proportion on each tissue. Horizontal line represents Bonferroni cutoff and only significant cell types are labeled.

## Notes

https://github.com/sudmantlab/gene_expression_aging

### Competing Interest Statement

The authors have declared no competing interest.

## Bibliography

1. B Charlesworth, Fisher, medawar, hamilton and the evolution of aging. Genetics 156, 927–931 (2000).

2. MR Rose, CL Rauser, G Benford, M Matos, LD Mueller, HAMILTONS FORCES OF NATURAL SELECTION AFTER FORTY YEARS. Evolution 61, 1265–1276 (2007).

3. A Viñuela, et al., Age-dependent changes in mean and variance of gene expression across tissues in a twin cohort. Hum. Mol. Genet. 27, 732–741 (2017).

4. B Balliu, et al., Genetic regulation of gene expression and splicing during a 10-year period of human aging. Genome Biol. 20 (2019).

5. M Somel, P Khaitovich, S Bahn, S Pääbo, M Lachmann, Gene expression becomes heterogeneous with age. Curr. Biol. 16, R359–R360 (2006).

6. S Wang, et al., Single-cell transcriptomic atlas of primate ovarian aging. Cell 180, 585–600.e19 (2020).

7. CP Martinez-Jimenez, et al., Aging increases cell-to-cell transcriptional variability upon immune stimulation. Science 355, 1433–1436 (2017).

8. C Cheng, M Kirkpatrick, Molecular evolution and the decline of purifying selection with age. Nat. Commun. 12 (2021).

9. K Jia, C Cui, Y Gao, Y Zhou, Q Cui, An analysis of aging-related genes derived from the genotype-tissue expression project (GTEx). Cell Death Discov. 4 (2018).

10. Genetic effects on gene expression across human tissues. Nature 550, 204–213 (2017).

11. O Stegle, L Parts, R Durbin, J Winn, A bayesian framework to account for complex nongenetic factors in gene expression levels greatly increases power in eQTL studies. PLoS Comput. Biol. 6, e1000770 (2010).

12. PH Sudmant, MS Alexis, CB Burge, Meta-analysis of RNA-seq expression data across species, tissues and studies. Genome Biol. 16 (2015).

13. P Sen, PP Shah, R Nativio, SL Berger, Epigenetic mechanisms of longevity and aging. Cell 166, 822–839 (2016).

14. ER Gamazon, et al., A gene-based association method for mapping traits using reference transcriptome data. Nat. Genet. 47, 1091–1098 (2015).

15. J Yang, SH Lee, ME Goddard, PM Visscher, Gcta: A tool for genome-wide complex trait analysis. The Am. J. Hum. Genet. 88, 76–82 (2011).

16. AM Newman, et al., Determining cell type abundance and expression from bulk tissues with digital cytometry. Nat. Biotechnol. 37, 773–782 (2019).

17. A Subramanian, et al., Gene set enrichment analysis: A knowledge-based approach for interpreting genome-wide expression profiles. Proc. Natl. Acad. Sci. 102, 15545–15550 (2005).

18. S Rath, et al., Mitocarta3.0: an updated mitochondrial proteome now with sub-organelle localization and pathway annotations. Nucleic Acids Res. 49, D1541–D1547 (2020).

19. R Cui, et al., Relaxed selection limits lifespan by increasing mutation load. Cell 178, 385–399.e20 (2019).

20. M Lek, et al., Analysis of protein-coding genetic variation in 60,706 humans. Nature 536, 285–291 (2016).

21. M Gayà-Vidal, M Albà, Uncovering adaptive evolution in the human lineage. BMC Genomics 15, 599 (2014).

22. A Liberzon, et al., The molecular signatures database hallmark gene set collection. Cell Syst. 1, 417–425 (2015).

23. AC Society, Cancer facts & figures (2022).

24. JG Tate, et al., COSMIC: the Catalogue Of Somatic Mutations In Cancer. Nucleic Acids Res. 47, D941–D947 (2018).

25. N Almanzar, et al., A single-cell transcriptomic atlas characterizes ageing tissues in the mouse. Nature 583, 590–595 (2020).

26. P Cheung, et al., Single-cell chromatin modification profiling reveals increased epigenetic variations with aging. Cell 173, 1385–1397.e14 (2018).

27. S Srivastava, The mitochondrial basis of aging and age-related disorders. Genes 8, 398 (2017).

28. S Tahmasebi, A Khoutorsky, MB Mathews, N Sonenberg, Translation deregulation in human disease. Nat. Rev. Mol. Cell Biol. 19, 791–807 (2018).

29. H Mostafavi, et al., Variable prediction accuracy of polygenic scores within an ancestry group. eLife 9 (2020).

30. C Giambartolomei, et. al, Bayesian test for colocalisation between pairs of genetic association studies using summary statistics. PLOS Genet. (2014).

31. M Wainberg, et al., Opportunities and challenges for transcriptome-wide association studies. Nat. Genet. 51, 592–599 (2019).

32. EE Porcu, et al., Mendelian randomization integrating gwas and eqtl data reveals genetic determinants of complex and clinical traits. Nat. Commun. 10 (2019).

33. TG Richardson, G Hemani, TR Gaunt, CL Relton, G Davey Smith, A transcriptome-wide mendelian randomization study to uncover tissue-dependent regulatory mechanisms across the human phenome. Nat. communications 11, 1–11 (2020).

34. MKR Donovan, A D’Antonio-Chronowska, M D’Antonio, KA Frazer, Cellular deconvolution of GTEx tissues powers discovery of disease and cell-type associated regulatory variants. Nat. Commun. 11 (2020).

35. JH Friedman, T Hastie, R Tibshirani, Regularization paths for generalized linear models via coordinate descent. J. Stat. Softw. 33, 122 (2010).

36. I Yanai, et al., Genome-wide midrange transcription profiles reveal expression level relationships in human tissue specification. Bioinformatics 21, 650–659 (2004).

